# Control of Modular Tissue Flows Shaping the Embryo in Avian Gastrulation

**DOI:** 10.1101/2024.07.04.601785

**Authors:** Guillermo Serrano Nájera, Alex M. Plum, Ben Steventon, Cornelis J. Weijer, Mattia Serra

## Abstract

Avian gastrulation requires coordinated flows of thousands of cells to form the body plan. We quantified these flows using their fundamental kinematic units: one attractor and two repellers constituting its Dynamic Morphoskeleton (DM). We have also elucidated the mechanistic origin of the attractor, marking the primitive streak (PS), and controlled its shape, inducing gastrulation flows in the chick embryo that are typical of other vertebrates. However, the origins of repellers and dynamic embryo shape remain unclear. Here, we address these questions using active matter physics and experiments. Repeller 1, separating the embryo proper (EP) from extraembryonic (EE) tissues, arises from the tug-of-war between EE epiboly and EP isotropic myosin-induced active stress. Repeller 2, bisecting the anterior and posterior PS and associated with embryo shape change, arises from anisotropic myosin-induced active intercalation in the mesendoderm. Combining mechanical confinement with inhibition of mesendoderm induction, we eliminated either one or both repellers, as predicted by our model. Our results reveal a remarkable modularity of avian gastrulation flows delineated by the DM, uncovering the mechanistic roles of EE epiboly, EP active constriction, mesendoderm intercalation and ingression. These findings offer a new perspective for deconstructing morphogenetic flows, uncovering their modular origin, and aiding synthetic morphogenesis.

## Introduction

Morphogenesis requires spatiotemporal coordination of hundreds of thousands of cells to sculpt a developing embryo [1, 2]. Elucidating the principles of morphogenesis, spanning mechanisms from molecular to organismal scales, is a grand challenge for the physics of living systems [3–7]. Tackling this challenge requires experimental advances and theoretical progress on two fronts: *i)* developing predictive biophysical models to test hypotheses and *ii)* devising mathematical methods to extract key insights from experimental data. Models of tissue flows include statistical and continuum theories for active matter [8], vertex models [9–11], and constitutive laws [12]. Recent work has also began to explore the interplay of signaling, mechanics, and geometrical constraints [1, 5, 13–21]. Meanwhile, methods to quantify tissue flows include statistical tools based on the connectivity between neighboring sites [22], and others quantifying cell shape changes and intercalation by mapping the temporal evolution of strain rates between neighboring cells [15, 23–25]. Yet, progress in *i* and *ii* have largely remained independent.

Recent imaging advances enable tracking of single-cell trajectories (**x**_*i*_(*t*)) in specific embryonic regions [26, 27]. In contrast, organism-scale data typically includes spatiotemporal velocities (**v**(**x**, *t*)) inferred from the collective motion of cells over whole tissues [28–30]. However, analyzing noisy datasets of **x**_*i*_(*t*) or **v**(**x**, *t*) over evolving domains can be overwhelming rather than informative. Ideally, one should discard unnecessary information and extract essential kinematic units robust to noise and invariant to the reference frames used to describe cell motion. This challenge, initially conceived to understand transport in passive fluids, led to the development of Lagrangian and Eulerian Coherent Structures [31–33], now widely adopted in science and engineering [34–36]. In morphogenesis, the Dynamic Morphoskeleton (DM) identifies dynamic ‘attractors’ and ‘repellers’—regions where cells converge or separate over a given time interval [37]—as fundamental kinematic units. The DM in chick [37–39], fruit fly [37], and zebrafish [26] embryos revealed early footprints of known morphogenetic features and new ones. Repellers, in particular, were previously undocumented despite their relevance for morphogenesis-mediated cell differentiation across organisms [26, 40]. Beyond kinematics, elucidating the mechanisms generating the DM provides a novel perspective to guide experiments and deconstruct morphogenesis in terms of distinct modules with physical, developmental and evolutionary implications [38, 39, 41, 42].

We focus on avian gastrulation, a crucial process in early development during which a flat sheet of approximately 60,000 cells breaks symmetry, setting the vertebrate body axes and forming the three germ layers (endoderm, mesoderm, and ectoderm). At the onset of gastrulation (HH1), avian embryos consist of a circular monolayer of suspended epithelial cells, the embryo proper (EP), surrounded by an annulus of extraembryonic tissue (EE) [43–45] (Fig. 1A). Initially, mesendoderm precursor cells are in a sickle-shaped region in the posterior EP. Over the next 15 hours, these mesendodermal progenitors converge and extend along the midline, undergoing individual cell ingression and forming the primitive streak (PS). This convergent extension is driven by active intercalation, apical constriction, and ingression [46–49] (Fig. 1A). These processes generate complex, embryo-scale coordinated cell movements and macroscopic vortical flows in the EP (Fig. S5). Simultaneously, the EE expands outward, powered by the active edge cell crawling on the vitelline membrane (Fig. 1A). The DM compresses the complex cell movements during avian gastrulation into one line attractor and two repellers (Fig. 1B, [37]). The attractor marks the PS, and its domain of attraction (DOA) identifies the sickle-shaped region of mesendoderm cells that will ingress into the PS. Repeller 1 (R1) partitions the EP and EE regions, while Repeller 2 (R2) bisects the DOA, segregating anterior and posterior mesendoderm cells.

**Figure 1:**
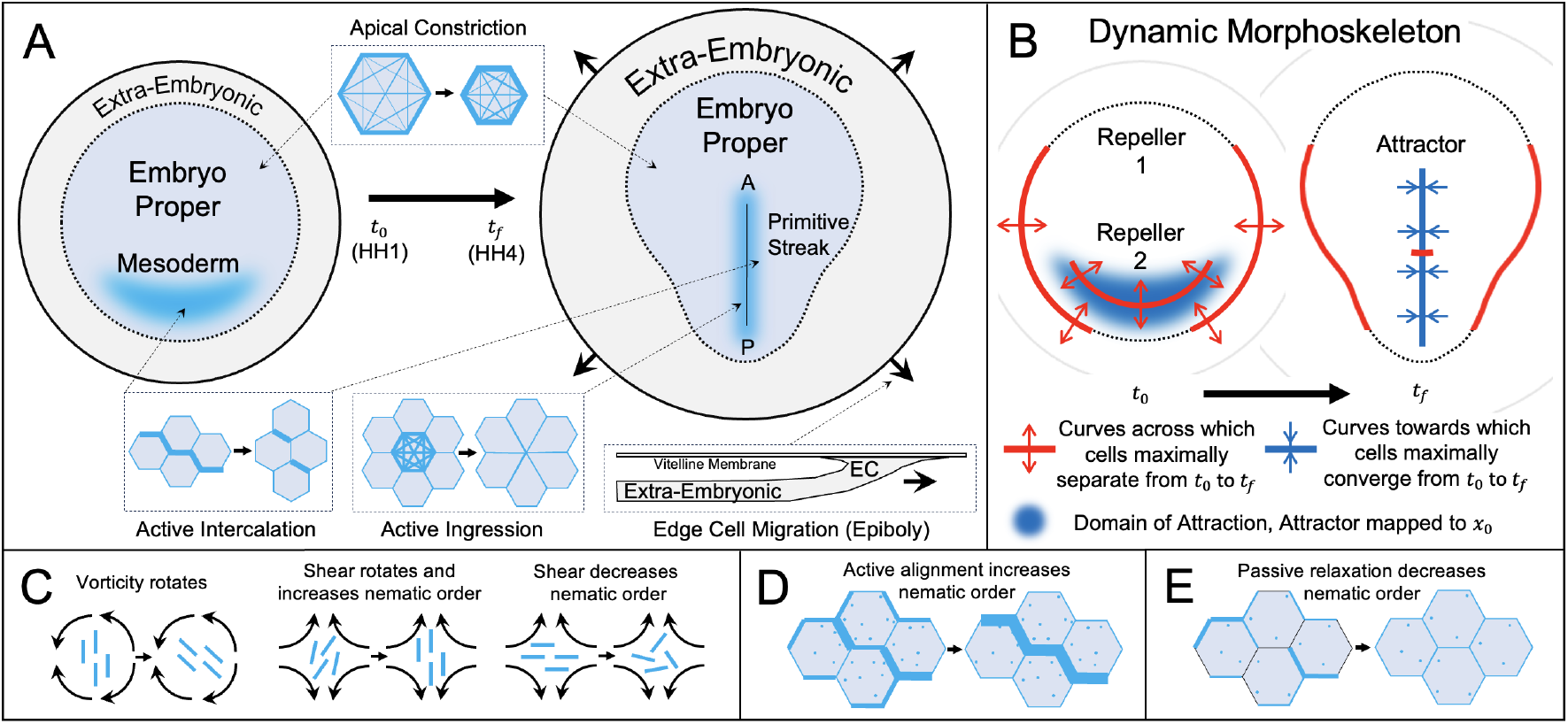
Avian Gastrulation. A) Avian morphogenesis involves extraembryonic (EE) expansion driven by edge cells (EC) crawling on the vitelline membrane, convergent extension of the presumptive mesendoderm to form the primitive streak (PS), and shape change of the embryo proper (EP). Myosin activity drives isotropic apical constriction throughout the EP and directs mesendoderm intercalations. Mesendoderm cells actively ingress at the PS, marking the anterior-posterior axis. Cyan in insets marks elevated myosin activity. B) The Dynamic Morphoskeleton of avian gastrulation. Repellers (R1 and R2) mark initial (*t*_0_) locations of cells that maximally separate by *t*_*f*_. The attractor marks final (*t*_*f*_) locations where cells maximally converge during [*t*_0_, *t*_*f*_]. The Domain of Attraction (DOA) marks the initial cell positions that converge to the Attractor. R1 lies on the EP-EE boundary, R2 bisects the presumptive mesendoderm, and the Attractor marks the PS. C-E) Active nematic tensor dynamics (Eq. (1c)). C) Flow coupling: Actomyosin cables (cyan nematic elements) reorient and align passively with the local flow (black). D) Actomyosin cables align active junctions through mechanosensitive myosin redistribution. E) Nematic order passively relaxes without activity or flow.

We previously developed a minimal 2D continuum model coupling active stresses and tissue motion, finding that gastrulation flows arise from a mechanosensitive active stress instability (Fig. S2A), confirmed by experiments [39]. This model demonstrated that changing the initial mesendoderm pattern and a parameter associated with active ingression can reshape the Attractor. Strikingly, model predictions matched experiments, showing that the Attractor of chick gastrulation flows can be altered to recapitulate other vertebrate gastrulation modes [38, 39, 42]. These findings convey two central messages: *i)* Mesendoderm shape and ingression capacity are sufficient control parameters to shape the Attractor and, therefore, the gastrulation mode. *ii)* Attractors compress **v**(**x**, *t*) into a robust, objective, Lagrangian (i.e., integrated over cell paths) kinematic unit with biological significance. The insights enabled by this compact representation would be inaccessible by comparing spatiotemporal **v**(**x**, *t*) among phenotypes [37, 39].

In this study, we investigate three open questions in avian gastrulation: the mechanistic origins of the morphogenetic repellers, their modularity, and the mechanisms controlling the dynamic embryo geometry. To address these questions, we develop a compressible active nematic model, building on our recent work [39]. This model predicts the emergence of R1 and R2 from distinct mechanisms and suggests their independent manipulability (modularity). To test these predictions, we developed novel, model-inspired mechanical and molecular perturbations, showing that R1 and R2 can indeed be separately controlled, *in vivo*, in chick embryos. Remarkably, the model also predicts the EP shape change from a circular disk to a pear shape as a self-organizing process, a feature beyond the scope of existing continuum models [21, 39, 50]. Our findings suggest a new perspective: morphogenetic flows can be decomposed into modular kinematic units, identifiable by the DM from data, with relatively independent mechanistic origins and functions. This kinematic modularity, distinct from genetic [51,52] or mechanical [53] modularity, provides a powerful lens for elucidating and controlling self-organization in natural and synthetic active matter.

## Mathematical Model

During avian gastrulation, embryonic cells frequently exchange neighbors, effectively behaving as a viscoelastic or viscous, compressible fluid deformed by active forces [21, 39, 50, 54]. Actomyosin cables spanning 6-8 cells generate active stresses [49], driving directed cell intercalations [46–48]. We previously quantified the intensity and orientation of myosin activity [39] and modeled actomyosin cables as contractile nematic elements, consistent with findings that nematic order dominates in epithelia at large length scales [55]. In [39], myosin activity generates active stresses that drive tissue flows, altering active stress patterns. This continuum model is computationally fast, interpretable, and uses fewer parameters than more microscopic approaches [56, 57]. Using fixed parameters and no spatial or temporal fitting, the model predicts EP gastrulation flows, specifically their attractors in wild-type (WT) and perturbed embryos [39].

This model, however, has important limitations. First, it enforces a fixed circular EP with no EE tissue. These preclude predictions of embryo shape change [45] and repellers, which are prominent in avian morphogenesis (Fig. 1B). Second, it precludes modeling relevant distinct EP-EE dynamics. Third, it assumes no dynamics for the myosin activity’s nematic order parameter. Here, we explicitly account for the EE tissue, which immunostaining reveals to be largely devoid of myosin activity [39]. In addition to myosin activity *m* and its average orientation *ϕ*, we model its dynamic anisotropy with a nematic order parameter *s*, which modulates anisotropic active stress ∝ *m***Q**, where **Q** = *s/*2[cos 2*ϕ*, sin 2*ϕ*; sin 2*ϕ*, −cos 2*ϕ*] is the nematic tensor. The *s* dynamics depends on flow coupling, active cable formation, and passive relaxation, enabling the creation, evolution and destruction of actomyosin cables (Sec. S2.2), as observed in experiments. Equation (1) summarizes our model:

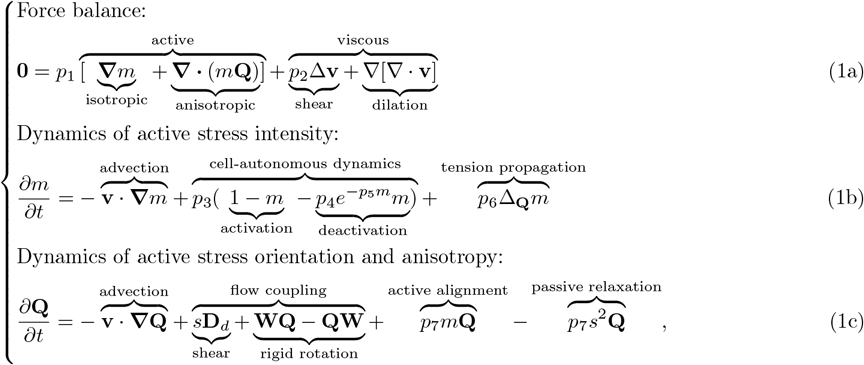

where **D**_*d*_ and **W** are the deviatoric strain rate and vorticity tensors, and *p*_*i*_ are non-dimensional parameters (Sec. S2.5). In Eq. (1a), *p*_1_ is the ratio of active stress to the characteristic viscous bulk stress and *p*_2_ is the ratio of shear to bulk tissue viscosities. Eq. (1b) describes the dynamics of *m*, the fraction of locally available myosin generating active stresses in (1a) (Sec. S2.1). *p*_3_ is the ratio of gastrulation’s characteristic timescale (*t*_*c*_ ≈ 15 hours) to the cumulative timescale for converting all available myosin into tissue-scale active stress (Sec. S2.1). *p*_4_ is the ratio of myosin tissue-scale activation to deactivation timescales. *p*_5_ is the mechanosensitivity of the deactivation rate to cable tension [39]. *p*_6_ is the ratio of mechanical (or tension-induced) propagation of myosin activity along cables [57, 58], to transport via advection [39]. The first three terms in Eq. (1c), standard in active nematics [59–61], account for transport and flow-induced effects on **Q** (Fig. 1C). Notably, with only these terms, convergent extension invariably destroys initial cable alignment, driving *s* to 0 in the vicinity of the streak (Fig. S2C, Sec. S2.2), inconsistent with experiments (Fig. 1 of [39]). To sustain convergent extension, an alignment mechanism is needed to counteract order destruction from self-induced flows (Fig. S2D). The active alignment term (*p*_7_*m***Q**) models actomyosin cable formation as an active process, polarizing myosin activity by redistributing myosin to junctions aligned with the direction of highest active stress [48, 49, 62, 63] (Fig. 1D). This reflects in the continuum setting a mechanism recently explored in vertex models, with local tension feedback redistributing myosin to increase anisotropy while conserving *m* [57,64]. By contrast, passive relaxation (−*p*_7_*s*^2^**Q**) reduces *s* accounting for cell shape rigidity and random intercalations [64, 65] (Fig. 1E). *p*_7_ is *t*_*c*_ times the activity-induced alignment and passive relaxation rate (Sec. S2.5). Active alignment and passive relaxation terms, introduced in active nematics [59,66], are yet unexplored in the context of morphogenesis.

As boundary conditions in Eq. (1a), we prescribe an outward normal velocity *v*_*e*_ consistent with observed migrating edge cells (Sec. S2.6, Fig. S3). For Eqs. (1b-1c) we impose no flux for *m* and **Q**. For the initial condition of Eq. (1b), we use immunostaining data indicating minimal myosin activity in the interior EE throughout gastrulation, while myosin activity increases over time in the EP, consistent with an *m* instability (Fig. S2A, Sec. S2.1). To account for the distinct myosin dynamics of EP-EE cells, we set *m*(**x**_EP_, *t*_0_) *> m*^*^ and *m*(**x**_EE_, *t*_0_) *< m*^*^ where *m*^*^ is the unstable uniform fixed point in the myosin dynamics (1b) (Fig. S2A). Additionally, we initialize higher myosin activity in the posterior EP, matching the sickle shape of mesendoderm precursors [38, 39] (cf. scalar field in Fig. 2A). For the initial condition of Eq. (1c), we set *ϕ*(**x**, *t*_0_) along the tangential direction, matching experimental observations [39], and determine the initial nematic order from *m*(**x**, *t*_0_): 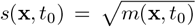, the equilibrium of Eq. (1c) before the onset of flows (Fig. S2B, Sec. S2.2). This reflects the expectation that a mild tangential cable alignment is amplified by active contraction and myosin redistribution in the mesendoderm precursor region (Fig. 2A) in line with experiments [39, 48]. For details on boundary and initial conditions and the numerical scheme, see Sec. S2.6-S2.8.

**Figure 2:**
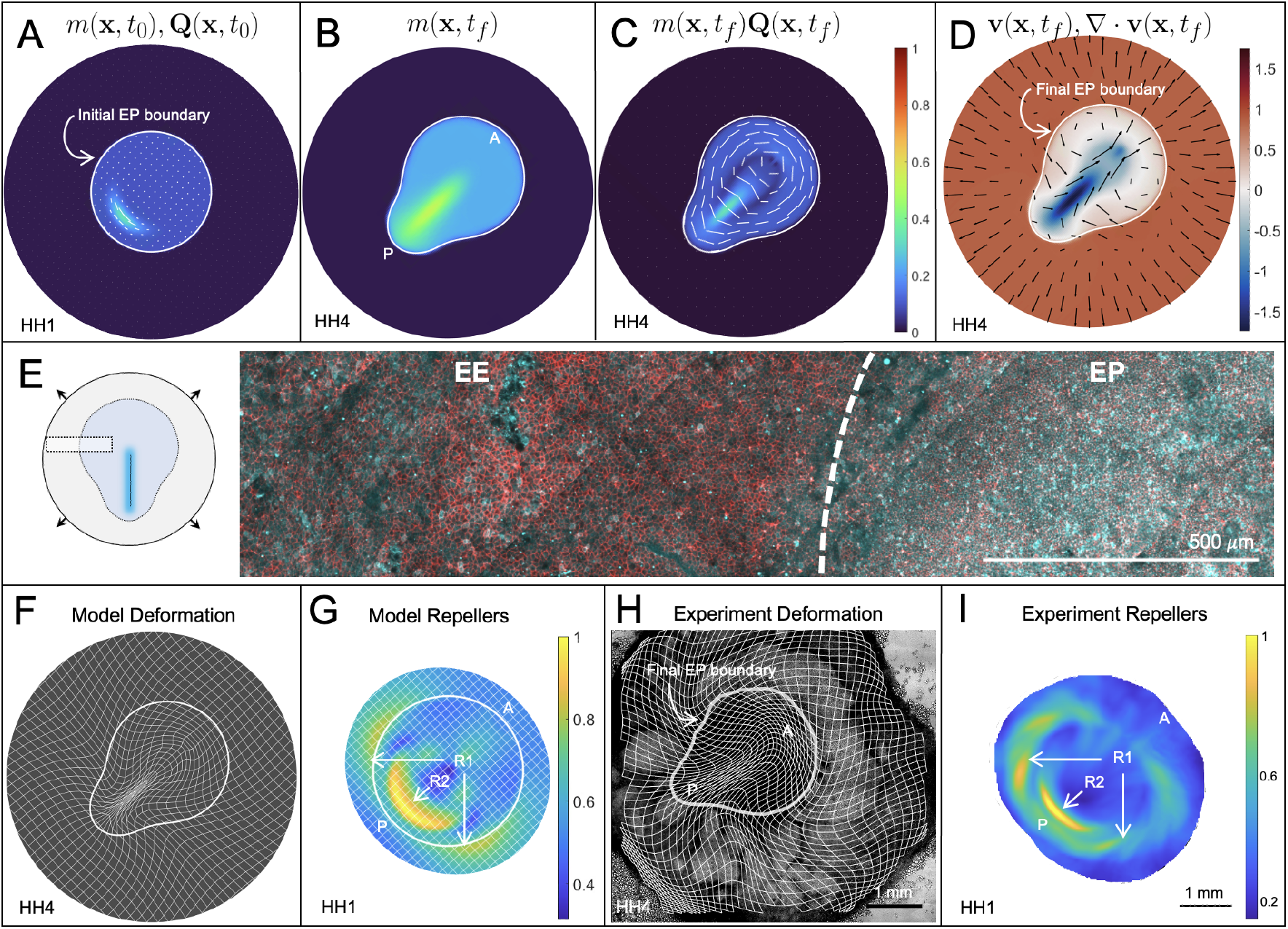
Wild-Type Gastrulation. A) Initial distribution of myosin activity *m* with orientation *ϕ* (white director field). Director lengths increase with *s* (exponentially to ease visualization). White circle (A, G) marks the initial EP-EE boundary, updated at later times (B-D, F) by advection with the model **v**. B,C) *t*_*f*_ distribution of isotropic (*m*) and anisotropic (*ms* with *ϕ, s* as in A) myosin activity. A-C share the color bar. D) Predicted *t*_*f*_ model velocity and divergence. All model quantities are dimensionless. E) Confocal image of a representative section of an HH4 stage WT embryo at the position indicated on the left. Staining for actin (phalloidin, red) and doubly phosphorylated myosin light chain (cyan) in the embryo. F) Model deformed Lagrangian grid and final EP-EE boundary. G) Model repellers (R1 and R2), marked by ridges of 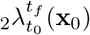 normalized by the spatial maximum, and displayed on the initial undeformed configuration (Sec. S1). H) Experimental grid overlaid on a bright-field image of an HH4 embryo. Final EP-EE boundary located in experiments drawing an initial polygon around R1 (Sec. S5.4) and advecting it with experimental **v**. I) Experiment repellers (R1 and R2), as in G. A, P labels mark the anterior-posterior axis. See Movie 1 for the WT model velocity, velocity divergence, isotropic and anisotropic stresses, active forces, repellers, attractors, and Lagrangian grids over time. Movie repellers and attractors colorbars are normalized over their spatiotemporal maxima, achieved at *t*_*f*_ = *HH*4.

## Results

### Model predicts repellers and embryo shape change

Solving Eq. (1), with the above initial and boundary conditions, we recapitulate avian gastrulation flows, including its characteristic Eulerian features: dynamic active stress patterns (Fig. 2A-C), vortices, and convergent-extension flows (Fig. 2D); and essential Lagrangian features: R1, R2, and EP shape change (Fig. 2F-G). See Movie 1 for the time evolution of these fields. Figure 2B and 2C show increasing EP myosin activity *m*(**x**, *t*_*f*_) and anisotropic activity increasing (magnitude of *m*(**x**, *t*_*f*_)**Q**(**x**, *t*_*f*_)), consistent with experiments [39]. Figure 2D shows velocity and velocity divergence at *t*_*f*_ = HH4 consistent with observations of sustained vortical flows, negative divergence in the PS (attractor), lower divergence magnitude in the rest of the EP and positive divergence in the EE region [39, 48] (Fig. S7). Immunostaining of HH4 embryos confirms high active myosin in the EP and low in the EE (Fig. 2E), consistent with our previous work [39], and in sharp contrast with reports in quail embryos [50], suggesting the presence of a tensile ring caused by a localized high *m* at the EP-EE boundary.

To assess Lagrangian features, Fig. 2F shows an initially uniform grid and circular EP-EE boundary (Fig. 2A) advected with the model **v**, revealing lower expansion of the EP relative to the substantial stretching of the EE [58]. With the EP-EE boundary now free to deform, the model also successfully predicts the EP’s long-recognized geometric transformation from circular to pear-shaped [45, 67] (also plotted in Fig. 2B-F). This nontrivial aspect requires the correct self-organizing dynamics of active forces (Movie 1 and Sec. S3.1) on the EP boundary, internal points of our modeling domain. Notably, other continuum models have been unable to reproduce EP shape change [39, 50, 68]. Figure 2G shows the highest Lagrangian stretching field 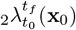, displayed at the embryo’s initial configuration, from which we extract the repellers [37]. Throughout, 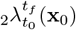 denotes the largest singular value field of the Lagrangian deformation gradient tensor 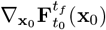, where 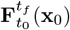 denotes trajectories from *t*_0_ to *t*_*f*_ starting at **x**_0_ (Sec. S1). High values of 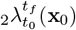 mark initial embryo locations (repellers R1 and R2) where nearby cells maximally separate by *t*_*f*_ (Fig. S1). To visualize these repellers (constituting the DM), we normalize 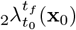 by its maximum. The model reproduces the circular arcs of R1 at the EP-EE boundary and R2 in the posterior [37] (c.f. Fig. 1B). Altogether, these results showcase the model’s strong predictive capability, providing only *m*(**x**, *t*_0_) and *ϕ*(**x**, *t*_0_) and scalar parameters as inputs.

To test our model, we developed an imaging and analysis pipeline using bright-field microscopy, enabling us to simultaneously track tissue flows in up to ∼16 embryos without fluorescent labels (Fig. S5). This approach revealed the robustness of the DM across several embryos, despite the intrinsic variability of their tissue flows (Movie S12 and Fig. S10). The DM remains tractable and reveals conserved features of the morphogenetic flows, even when Eulerian approaches struggle to capture meaningful patterns [26, 37]. The DM’s accessibility and robustness provide a strong baseline for interpreting perturbations, facilitating high-throughput screens to identify novel factors influencing the DM. Representative Lagrangian deformation (Fig. 2H) and repellers (Fig. 2I) from this pipeline are consistent with model predictions (Fig. 2F-G). Importantly, our goal is not to precisely match the DM between model and experiment by fitting parameters, as this would depend on specific experiments (Fig. S10). Instead, we aim to predict the existence and average geometry of the repellers, which are remarkably robust across experiments (Fig. S10).

### Origin and elimination of Repeller 1

We next investigate the mechanisms underlying R1. Edge cells (ECs) adhering to the VM crawl outwards, pulling the EE and EP radially (Fig. 1A). EC crawling rates are nearly constant and robust to several treatments, including excision of the EP [69–71], justifying our boundary condition for **v** (Sec. S2.6). Epiboly induces global tension [68, 72–74] that propagates to the EP, as severing the EP-EE boundary causes both regions to contract [73]. Yet, R1 shows that EP and EE cells separate, implying distinct expansion rates. We hypothesize that EP resists epiboly-driven expansion with isotropic actomyosin activity (Figs. 1A, 2A,B,E), which constricts cells’ cortical cytoskeletons, allowing them to bear the isotropic tension imposed by epiboly [73] without stretching like EE cells [68]. Isotropic myosin activity can constrict cells via circumferential purse-string contraction [75] or medial apical network contraction [76, 77] (Fig. 1A). Before the onset of motion, apical areas in the EP are already slightly smaller in the embryo and smallest in the posterior [48, 78], consistent with elevated myosin activity. As gastrulation proceeds, EP tissue thickens, and apical cell areas decrease [74, 78]. Decreasing EP cell areas accompany the marked rise in EP myosin activity, with the greatest shrinkage near the PS where myosin activity grows highest [48, 78]. In contrast, EE cells, devoid of active myosin (Fig. 2E), stretch thin, and their cell areas can increase more than double [68, 78]. These distinct EP-EE dynamics are consistent with the model’s patterns of myosin and cumulative deformation (Figs. 2A,B,F).

To test our hypothesis that R1 results from the opposition between EP inward constriction and EC outward crawling, we first eliminated epiboly in the model, solving Eq. (1) with the same parameters, initial and boundary conditions as in Fig. 2, except *v*_*e*_ = 0. Eliminating epiboly dramatically weakens differential EP-EE expansion and effectively eliminates R1 relative to R2 and EP shape change, which are not substantially altered (Figs. 3A-B). Eliminating isotropic myosin activity or the distinct EP-EE *m*(**x**, *t*_0_) is also sufficient to eliminate R1 in the model (Sec. S3), but these perturbations are experimentally impractical. Additionally, eliminating epiboly may affect the average isotropic tension of the tissue [74], reducing the rate of myosin accumulation, but any such effect is beyond the scope of our current model. To experimentally verify our prediction, we restricted epiboly movements. Past approaches to interfering with epiboly have limitations: ablating ECs blocks epiboly only transiently as new ECs quickly differentiate and resume epiboly [72]; removing the EE entirely can compromise development because of its signaling role patterning the early EP [73, 79]; chemical treatments like Colchicine also disrupt cell division [80, 81]; and manually wrinkling the VM [73] or extracting yolk [82] to reduce VM tension only partially reduce expansion and not always uniformly. To overcome these limitations, we developed a novel technique, cauterizing the inner face of the VM surrounding the EE using a soldering iron *ex vivo* (Fig. 3C and Sec. S5.2). The ECs cannot adhere to the cauterized region, creating a fixed boundary that experimentally realizes *v*_*e*_ = 0 uniformly.

**Figure 3:**
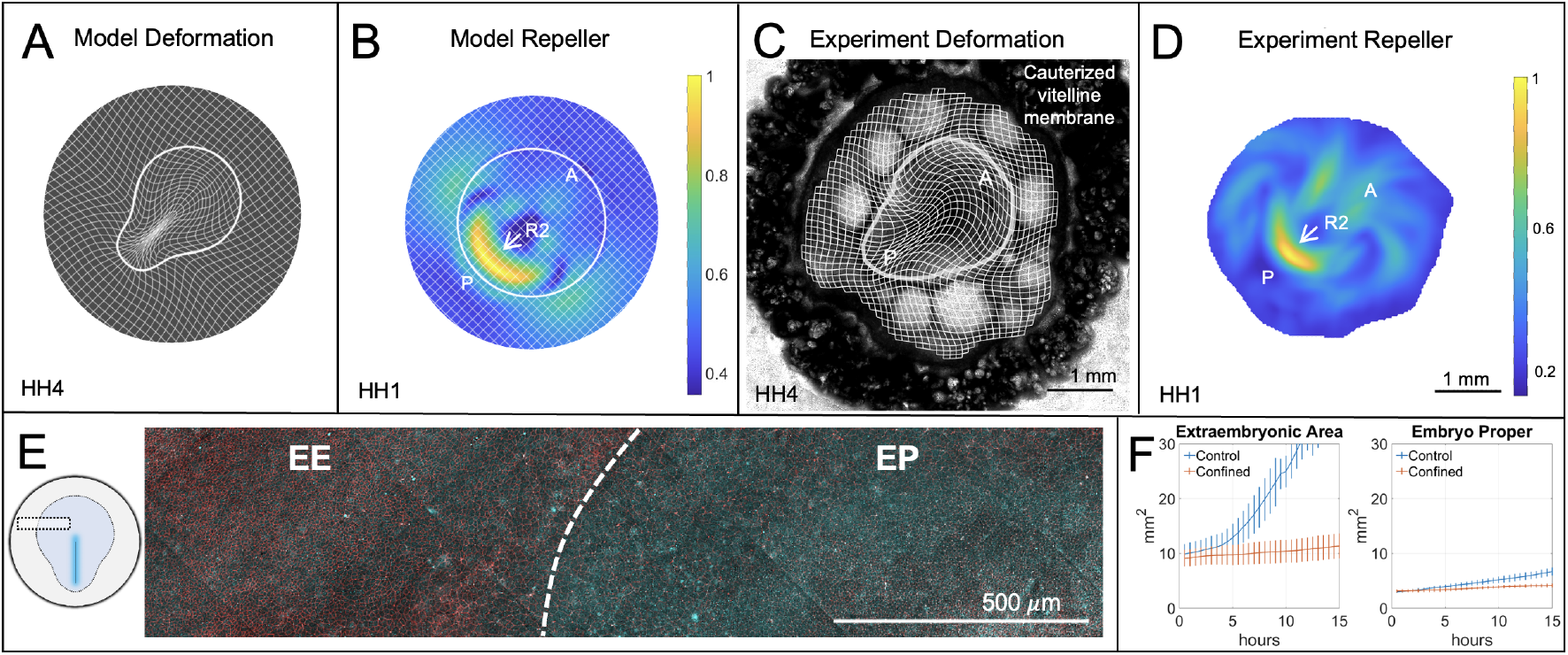
Repeller 1 Elimination. Confining the embryo eliminates R1 but preserves R2. A,B) Model results solving Eq. (1) with *v*_*e*_ = 0 reflecting embryo confinement and all other parameters as in WT. A) Deformed Lagrangian grid and final EP-EE boundary. B) Model repeller (R2), marked by a ridge of 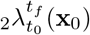, normalized by the spatial maximum, and displayed on the initial undeformed configuration. C,D) Embryo confined in experiments by cauterizing the EP side of the vitelline membrane. C) Experimental Lagrangian grid overlaid on the bright-field image of a confined HH4 embryo, boundary located as in Fig. 2. D) Experiment repeller (R2), as in B. Compare A-D with the corresponding WT Figs. 2F-I. A,P labels mark the anterior-posterior axis. Movie 2 shows the time evolution of Lagrangian and Eulerian model fields as in Movie 1. E) Doubly phosphorylated myosin light chain (cyan) and actin (phalloidin, red) in a representative section of a confined embryo. F) Mean and standard deviation of experimental EP and EE areas over 15 hours (*N* = 8 per group).

This confinement technique is compatible with our high-throughput live imaging pipeline (Fig. S5). A representative Lagrangian deformation grid of the confined embryo (Fig. 3C) shows lesser differential EP-EE stretching than WT (Fig. 2H) and that the embryo still becomes pear-shaped (Fig. 3C) and forms a PS (Figs. S6B and S9B). Figure 3D indicates the presence of R2 but no R1 (compare with Fig. 2I), consistent with model predictions (Fig. 3B). Immunostaining of phosphorylated (active) myosin verifies persistent contractility across the EP under confined conditions (Fig. 3E and Fig. S8). Figure 3F confirms that confinement prevents embryo expansion and that, despite persistent EP contractility, the marked difference in EE-EP expansion observed in the WT embryo (blue curves) has been reduced (orange curves). Remarkably, we find that the confined embryos develop well past gastrulation stages in these conditions (Fig. S9), suggesting that epiboly (and therefore R1) may be unnecessary at gastrulation stages (Discussion and Sec. S4). These results suggest that R1 arises from a tug-of-war between EE epiboly and EP active constriction. Without epiboly, no substantial separation occurs at the EP-EE boundary.

### Origin and elimination of Repeller 2

Preserving R2 without R1 suggests that avian gastrulation’s repellers may arise from independent mechanisms. We hypothesize that R2, bisecting the initial crescent-shaped mesendoderm, results from actomyosin cable-directed intercalations in this region [48,49], independent of epiboly and EP constriction associated with R1. To test our hypothesis, we eliminated the model mesendoderm, represented by elevated posterior myosin activity and anisotropy, solving Eq. (1) with the same parameters, initial and boundary conditions as in WT, but initializing uniform *m* in the EP. This also eliminates the initially elevated posterior cable anisotropy associated with active alignment (Fig. 2A, Sec. S2.7). With this change, the EP remains circular and retains differential EP-EE expansion (Fig. 4A). Figure 4B shows the elimination of R2, but R1 remains and becomes circularly symmetric due to the retention of epiboly and uniform isotropic active stress throughout the EP. Eliminating anisotropic active stress or active cable formation (*p*_7_ = 0) is also sufficient to eliminate R2 (Sec. S3).

**Figure 4:**
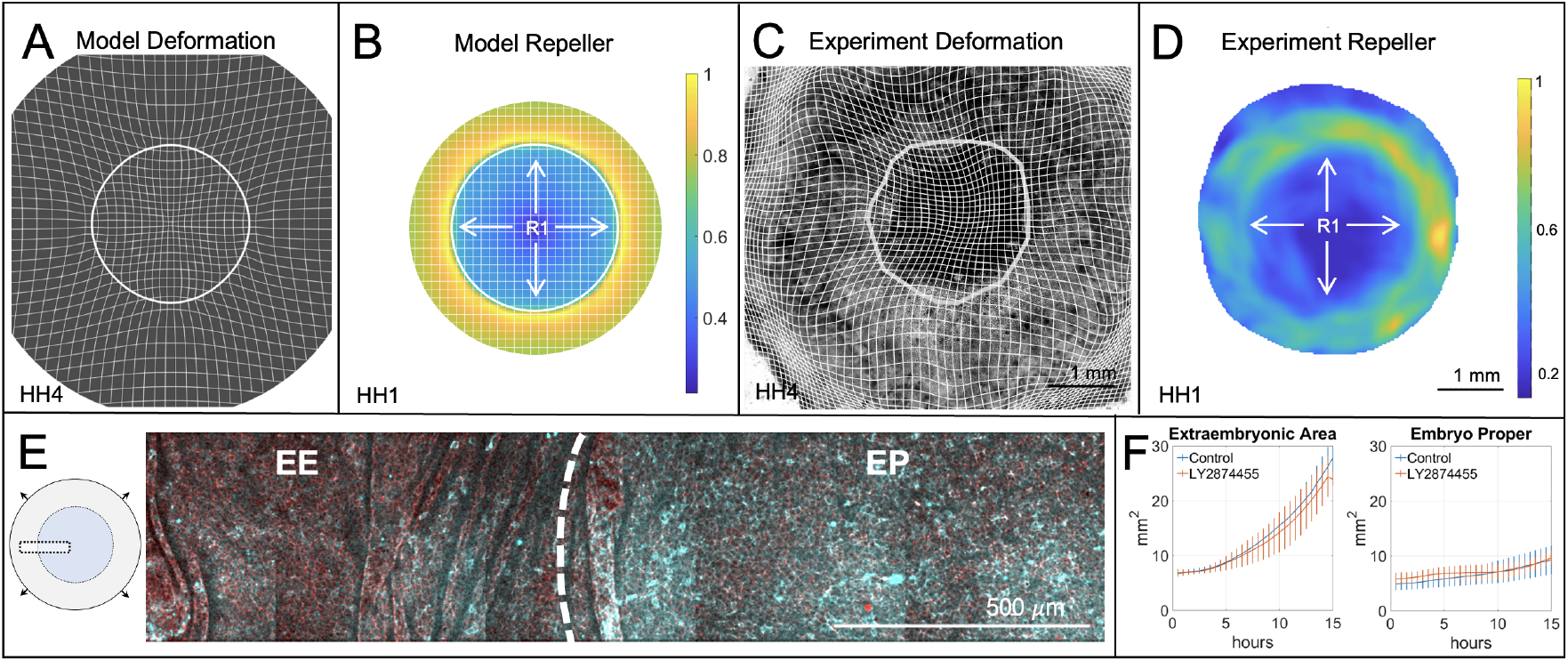
Repeller 2 Elimination. Blocking mesendoderm induction eliminates R2 but preserves R1. A,B) Model results obtained solving Eq. (1) with uniform *m*(**x**_*EP*_, *t*_0_) reflecting the absence of mesendoderm. A) Deformed Lagrangian grid and final EP-EE boundary. B) Model repeller (R1), marked by a ridge of 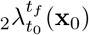, normalized by the spatial maximum, and displayed on the initial undeformed configuration. C,D) Mesendoderm induction blocked in experiments using 1*µM* LY2874455, a pan-FGF receptor inhibitor. C) Experimental grid overlaid on bright-field image of a confined HH4 embryo, boundary located as in Fig. 2. D) Experiment repeller (R2), as in B. Compare A-D with the corresponding WT Figs. 2F-I. Movie 3 shows the time evolution of model fields as in Movie 1. Movie attractor colorbar is normalized using the WT spatiotemporal maximum to emphasize the relative lack of deformation as in Fig. S6. E) Doubly phosphorylated myosin light chain (cyan) and actin (phalloidin, red) in a representative section of a treated embryo. F) Mean and standard deviation experimental EP and EE areas over 15 hours (*N* = 4 per group).

We previously showed that oriented cell intercalation and oriented myosin cables are characteristic properties of the mesendoderm cells [38]. Indeed, using a pan-FGF receptor inhibitor LY2874455 [83] completely blocks mesendoderm formation and the oriented cell intercalation associated with PS formation [37, 38]. To verify our model prediction, we experimentally blocked mesendoderm formation using the same chemical treatment (LY2874455 1*µM*) and live imaged whole gastrulas to obtain PIV **v**. Consistent with model results, the embryo retains a circular geometry (Fig. 4C), keeping R1, which becomes circularly symmetric, while eliminating R2 (Fig. 4D) and the Attractor (Fig. S6). Immunostaining of treated embryos shows that elevated phosphorylated myosin in the EP is not affected by the FGF inhibition (Fig. 4E). Further, Fig. 4F demonstrates that blocking R2 leaves the EP and EE areal expansions unaltered compared to WT. These results strongly indicate that the elevated EP actomyosin activity represents an inherent property of the EP, not requiring mesendoderm induction. Instead, induction is only required to assemble anisotropic cables perpendicular to the midline and execute the directed intercalations that contribute to forming the PS (the attractor) and embryo shape change. Finally, combining embryo confinement with inhibition of mesendoderm induction (*v*_*e*_ = 0 and uniform EP *m*(**x**, *t*_0_)), we simultaneously eliminated both repellers in the model and experiments (Fig. 5A, last column). Without epiboly and mesoderm-driven active intercalation, minimal deformation occurs, resulting in no differential EP-EE expansion, shape change, or Attractor (Fig. S6).

**Figure 5:**
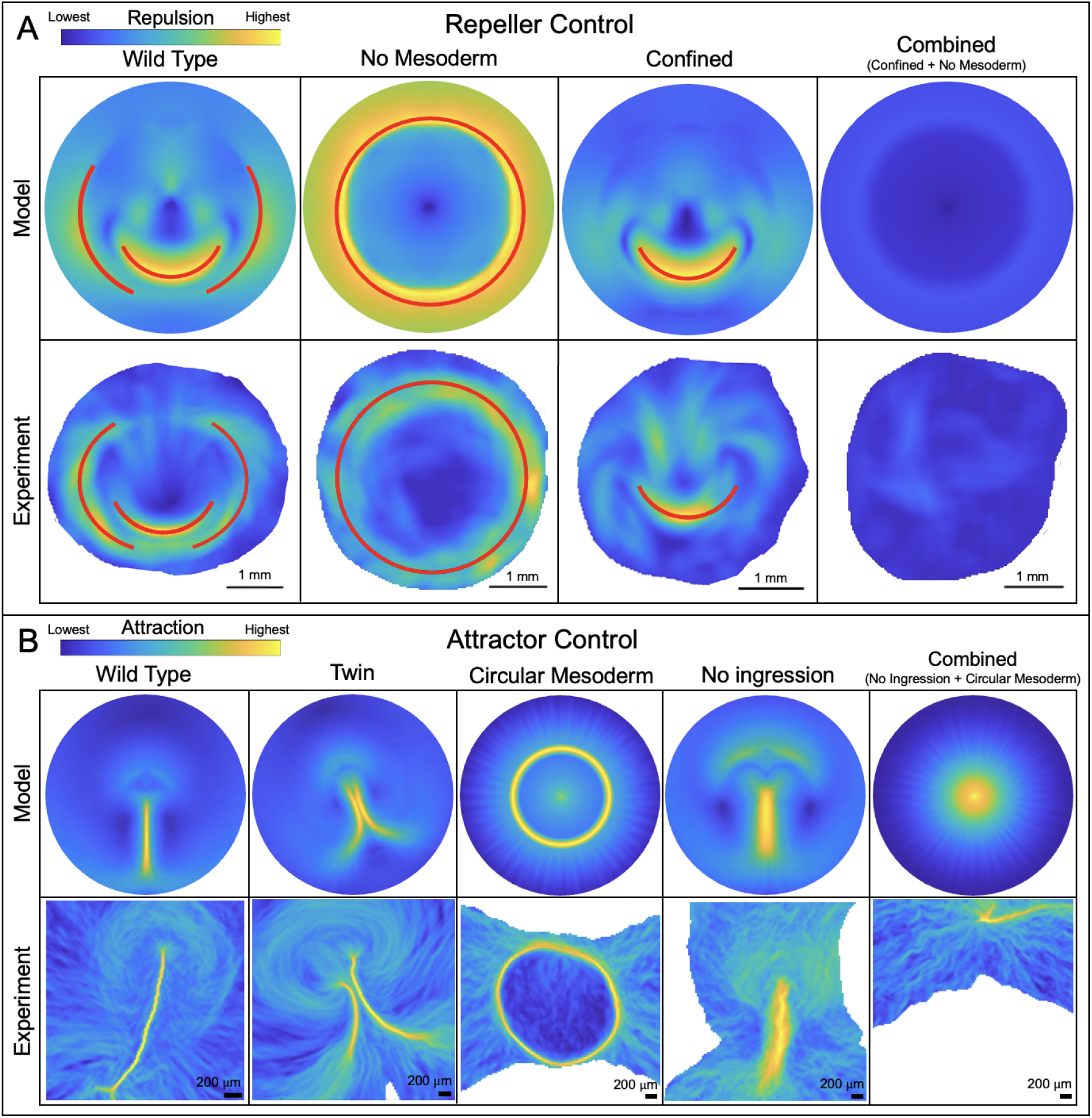
Controlling the Dynamic Morphoskeleton: Repellers and Attractors. A) Repellers R1 and R2 can be controlled combinatorially by confining the embryo and inhibiting mesendoderm induction. First three columns adapted from Figs. 2-4. The fourth column shows the perturbation obtained by simultaneously confining the embryo and inhibiting mesendoderm induction in the model and experiments. Repellers for the combined perturbation are normalized by the WT spatial maximum to emphasize the lack of deformation. Movie 4 shows the time evolution of model fields associated with the fourth column, normalizing the repeller and attractor fields by the WT spatiotemporal maxima. All other fields of 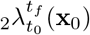 are normalized by their spatial maxima with repellers marked in red. B) Control of attractor shape in model and experiments by modulating cells’ ability to ingress and initial mesendoderm shape. Fields show 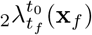, normalized by their spatial maximum. Panels adapted from our previous work [38, 39].

## Discussion

In this study, we have identified the mechanisms and cellular behaviors underlying multicellular flow repellers during avian gastrulation and how embryos change geometry from circular to pear-shaped (Figs. 1-2). Combining in-vivo experiments, active matter theory and nonlinear dynamics, we found that R1, separating EE from EP, arises from the balance between outward epiboly and active constriction of EP cells (Fig. 4). In contrast, embryo shape changes, PS-formation (Attractor), and R2 require active intercalations by mesendoderm cells, sustained by active cable formation (Fig. 3). The circular symmetry of R1 in the absence of R2 (Fig. 4) clarifies their interaction in WT (Fig. 2). With both repellers present, the EP shape change associated with R2 accentuates EP-EE separation at the lateral posterior EP edges, breaking R1’s otherwise circular symmetry. The relative strength (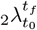 ridges) of R1 and R2 and the degree to which they interact may depend on the relative timing of epiboly and mesoderm-driven convergent extension (Fig. S4).

Our findings were guided by the DM, which compresses noisy spatiotemporal tissue flows into their fundamental kinematic units (attractors and repellers) (Fig. 1B) [37], robustly to intrinsic embryo variability (Fig. S10). These units guided model development and experiments to identify their originating mechanisms. The independent mechanisms underlying R1 and R2 and their manipulability suggest they are modular. This resonates with the notion of morphogenetic modularity: complex systems, like developing embryos, are composed of semi-autonomous modules, modifiable without disrupting the overall system’s functionality [53, 84]. The combinatorial elimination of R1 and R2 (Fig. 5A) has implications for avian morphogenesis control. In our recent findings, we also controlled avian gastrulation’s Attractor (Fig. 1B) [38, 39], modifying its WT line shape (the PS) into a ring, a dot, or a thicker shorter line (Fig. 5B) by independently modulating the initial mesendoderm precursor shape and cells’ capacity to ingress, recapitulating alternative gastrulation modes and providing evolutionary insights [42]. Modularity via DM helps understand the independent evolution of distinct aspects of development [84], and brings a fresh perspective to control natural and synthetic morphogenesis [41].

R1 arises from the tug-of-war between EE epibolic movements and the inherent contractility of the EP. Surprisingly, when we confined the embryo to eliminate epiboly while maintaining an intact EE, embryos still gastrulated and developed proportioned axial structures, despite having shorter body axes (Fig. S9). Similarly, a recent report showed that decreasing tension in the VM delayed epiboly, resulting in a shorter body axis [82]. Disrupting epiboly progression in zebrafish does not prevent gastrulation but also produces a shorter body axis [85, 86]. These findings suggest that while epiboly is crucial for nutritive functions later in development [87], it is not required for the early stages of avian embryogenesis. The contractility of the avian embryo may have evolved as a mechanism to resist the influence of epiboly, allowing the embryo to maintain its intrinsic patterning mechanisms and developmental timeline within the space delimited by R1. Indeed, experiments show that partial EP ablations normally require detachment of the EE from the VM to avoid the embryo ripping apart [88], suggesting that epiboly forces can pull apart a mechanically compromised EP. In contrast, R2 is required for primitive streak formation and embryo shape changes. The independent origins and modular nature of these repellers provide a new perspective on the robustness and evolvability of avian gastrulation (Sec. S4).

Our non-cell-autonomous model (Eq. (1)) consists of a viscous, compressible, active nematic flow driven by EE epiboly and EP-EE distinct myosin dynamics. *m* generates isotropic active stress modeling active cell constrictions and ingressions, and anisotropic ones generating active intercalations. Anisotropic active stresses arise from three key variables: cable orientation *ϕ, m* and order parameter *s*, quantifying the presence of aligned actomyosin cables, and whose dynamics involve non-standard activity-induced alignment and passive relaxation terms. These two processes capture the creation and destruction of aligned actomyosin cables, which are ubiquitous in experiments and critical for sustaining gastrulation flows. Using initial observed values of *ϕ, m* and scalar space- and time-independent parameters, our model accurately predicts 15 hours of avian gastrulation flows and embryo shape changes. This aspect, not addressed by existing models, is nontrivial as it requires correct prediction of active forces on the EP boundary, internal points of our modeling domain (Movies 1-4 and Sec. S3.1), dynamically shaping the embryo from a disc to a pear shape. Increasing shear viscosity, reflecting stiffer cell-cell junctions, results in poor EP shape change and disrupts vortical movements (Sec. S3). Theoretical [89–91] and experimental [50, 92, 93] studies suggest that division may contribute to tissue fluidity. In future work, we plan to *i)* incorporate divisions, ingressions and viscoelastic tissue rheology; *ii)* explore the biophysical role of a topological defect [60] in myosin anisotropy present in the anterior PS (Fig. 2C), colocated with the Hensen’s Node [94]; *iii)* compare our approach to the recent active solids description of morphogenesis [18, 64, 95].

In conclusion, our study combines in-vivo experiments, active matter physics and nonlinear dynamics to reveal how avian embryos control their shape and modular tissue flows driven by independent mechanisms. The DM facilitated these findings, shedding light on the robustness and evolvability of avian morphogenesis. Owing to its kinematic and compact nature, the DM can be instrumental in discovering new mechanisms controlling natural and synthetic morphogenesis, even when the underlying molecular mechanisms are not fully understood.

## Acknowledgements

The authors acknowledge Dillan Saunders for insightful discussions and Alexandra Neaverson, Yuri Takahashi, Apolline Delahaye and Sreejith Santhosh for their input and experimental support.

## Funding

GSN acknowledges support from Leverhulme Trust Early Career Fellowship (ECF-2022-474). A.P. acknowledges support from the National Institutes of Health (NIH) under training grant number T32-GM127235. CJW thanks the BBSRC (BB/N009789/1, BB/K00204X/1, BB/R000441/1, BB/T006781/1) for financial support and a Wellcome Trust imaging equipment award (101468/Z/13/Z) for partial support. MS acknowledges support from the Hellman Foundation.

### Author Contributions

G.S.N., C.J.W., and M.S. designed the research. G.S.N designed, performed, and analyzed experiments. A.P. and M.S. formulated the mathematical model. A.P. performed and analyzed numerical simulations. A.P., G.S.N., and M.S. wrote the manuscript. All authors contributed to the manuscript. M.S., C.J.W., and B.S. supervised the project.

## Supplementary Materials for

## S1 The Dynamic Morphoskeleton

The Dynamic Morphoskeleton (DM) locates dynamic attractors and repellers from tissue velocities [1] or cell trajectories [2]. Using only kinematic data, the DM is agnostic to the forces and microscopic mechanisms underlying coherent cell motion and can be computed over any developmental interval [*t*_0_, *t*] in the dataset. We denote the cell trajectories by

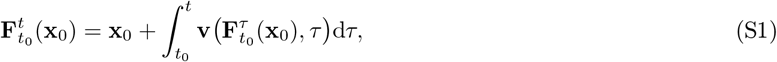

marking the time−*t* positions of cells that started at **x**_0_ at initial time *t*_0_ (Fig. S1A). The largest singular value 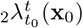 of the deformation gradient 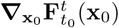 quantifies the maximum separation by time *t* between neighboring trajectories starting near **x**_0_ at *t*_0_

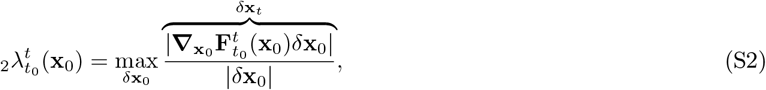

where *δ***x**_0_ parameterizes the neighborhood of **x**_0_. Repellers are identified from the ridges (i.e. regions of high values) of 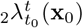, following trajectories forward in time (Fig. S1B). Likewise, attractors are computed from the flow map’s inverse, 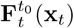, tracing trajectories back in time, and located by ridges of 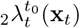 sdenoting the largest singular value of 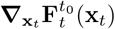. For more details and algorithm, see [1].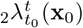 is related to the finite time Lyapunov exponent (FTLE) by 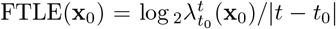. Taking the logarithm is appropriate in chaotic flows characterized by exponential stretching, but for slow morphogenetic flows, 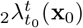 is more appropriate. Moreover, because we are interested in the presence or absence of the repellers and their average geometry, in the main text, we normalize 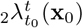 in each plot by its maximum spatial value. See Fig. S6 for absolute levels, exhibiting similar magnitudes of cumulative deformation in model and experiments. Fitting the experimental values precisely is less informative as the approximation of 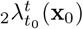 from experimental velocity data can be affected by its resolution [3], whereas the average geometry remains robust (Fig. S10).

**Supplementary Figure S1:**
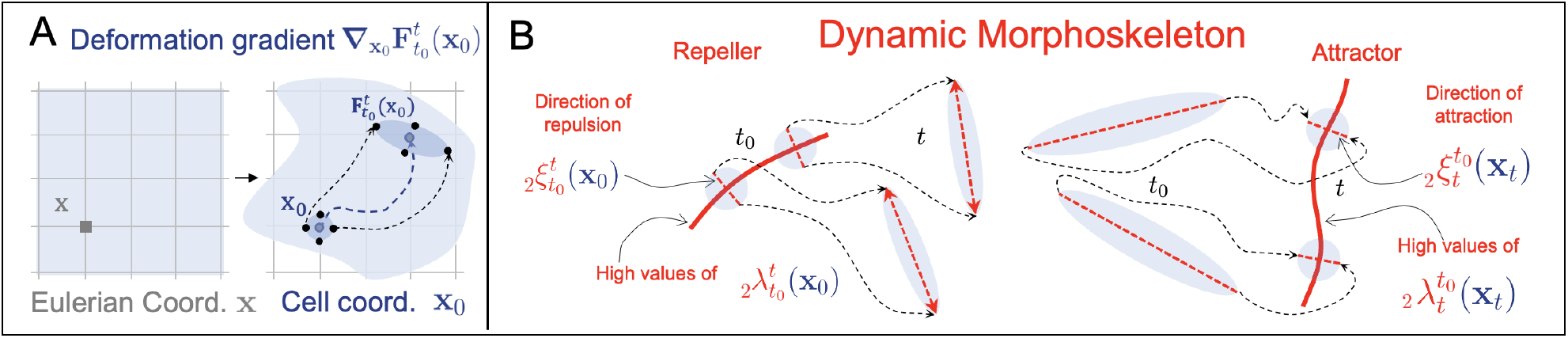
The Dynamic Morphoskeletons [1] Quantify Morphogenetic Flows. A) Eulerian coordinates **x** describe fixed spatial locations, while Lagrangian (cell) coordinates **x**_0_ label the identity of cells or tissue regions at their initial position and follow their trajectories 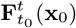. B) Fix the initial time *t*_0_ of the Lagrangian analysis, which we consider to be the beginning of our experiments (edevelopmental stage HH1). For any Lagrangian timescale *t* − *t*_0_, the Dynamic Morphoskeleton consists of repellers and attractors. Repellers mark regions at the initial cell configuration **x**_**0**_ across which cells maximally separate by time *t*. Attractors mark regions on the final tissue configuration towards which cells maximally converge by time *t*.

The DM is *i)* objective, i.e., invariant to time-dependent translations and rotations of the (arbitrary) coordinate system used to describe cell motion because based on tissue deformation, and *ii)* integrative, providing aggregate information along trajectories instead of isolated snapshots [1]. These properties make the DM robust to measurement errors, noisy data, and global artifacts such as embryo drifts. They also empower the DM to reveal dynamic patterns to which the velocity field and other non-objective or instantaneous metrics are blind [1, 2].

## S2 Mathematical Model

Avian gastrulation flows (planar velocity **v**(**x**, *t*) = [*u*(**x**, *t*), *v*(**x**, *t*)]) are driven by external boundary conditions (epiboly motion) and forces arising from gradients of active stresses. We model active stresses arising from myosin activity in a compressible Stokes flow, as in our previous work [4]. In the following sections, we recap our model and extend it in two key ways: *i)* Expand our domain to account for both the embryo proper (EP) and the surrounding extraembryonic (EE) tissue and model their distinct dynamics. *ii)* Model the dynamic anisotropy of active stress in addition to its intensity and average orientation. These extensions are necessary to understand the mechanistic origins of Repeller 1 (R1), Repeller 2 (R2) and the embryo’s dynamic geometry.

### S2.1 Active Stress Magnitude

Active stresses are generated by non-muscle myosin II (hereafter just myosin) contracting F-actin bundles at adherens junctions in the cells’ apical cortex [5, 6]. The amount of active myosin depends on *i)* the total myosin available for recruitment and activation and *ii)* the fraction of myosin activated. Since it is not well understood how *i)* changes during gastrulation, we assume that available myosin remains constant. Thus, increases in myosin activity correspond to increases in the fraction activated. We define *m*(**x**, *t*) as the fraction of available myosin active in a patch of tissue. Our previous work defined *m* in units of active stress, introducing the overall scale of myosin activity as an additional variable, controlled by the maximum available myosin concentration *m*_*n*_ [4]. Here, by defining *m* as a fraction of maximum available myosin, we eliminate this degree of freedom and instead convert myosin activity into isotropic and anisotropic stress with two constant parameters *α, β* (units of stress) defined in Section S2.3. This retains the same physics and is equivalent to non-dimensionalizing *m* in [4] using *m*_*n*_ and re-scaling other parameters accordingly.

The active stress magnitude, proportional to *m*(**x**, *t*), evolves as

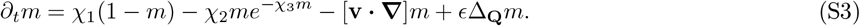

where *χ*_1_ and *χ*_2_ are effective rate parameters for the active stress dynamics due to activation and deactivation of myosin in a patch of tissue. Generating tissue-scale active stresses involves myosin binding to actin, phosphorylation, power strokes, and other mechanical processes [4, 6, 7]. The kinetics of these processes remain challenging to coarsen into tissue-scale models. In the chick embryo, successive, active intercalations can take up to an hour [6], consistent with other tissue-scale models in which myosin dynamics occur on the slowest timescale [8–10]. *χ*_1−3_ are chosen to ensure that *i)* the tissue-scale myosin dynamics preserve an active stress instability [4] (Fig. S2A) and *ii) m* does not saturate throughout the EP during gastrulation, consistent with experiments [4]. Total myosin availability could also exhibit heterogeneous dynamics associated with concurrent cell differentiation (i.e., mesendoderm cells may have higher availability). Yet, because such dynamics are not well characterized, we capture the key phenomenological features with Eq. (S3). *m* grows at a rate proportional to 1 − *m*, ceasing when all myosin is active (*m* = 1) and reaching its maximum rate when all myosin is inactive (*m* = 0). Myosin’s detachment rate instead exhibits mechanosensitivity, decreasing exponentially with tension [11], possibly due to a catch bond mechanism [12]. Because myosin activity largely determines junctional tension [13], the myosin deactivation rate decreases exponentially with *m*, with sensitivity *χ*_3_ as shown in [4]. The first three terms of Eq. (S3) represent average cell-autonomous dynamics in the Lagrangian (or cell) frame. Examining their spatially uniform fixed points reveals an instability (*m*^*^) between two stable equilibria [4]: a lower equilibrium corresponding to low baseline myosin activity and a higher equilibrium near saturation (≈ 1) (Fig. S2A). The last te(r[m accou]nts fo)r the directed propagation of active stress intensity along actomyosin cables [4]. We define 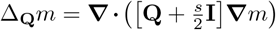 to represent directional tension propagation proportional to the degree of nematic order (our proxy for the presence of cables, see Section S2.2). For example, if *ϕ* = 0 (along *x*), **∇**_**Q**_*m* = ([*s* 0; 0 0] **∇***m*), ensuring propagation only in *x. ϵ* is the ratio between tension-propagated myosin activity and transport via advection, which we take to be small, as explained in [4].

### S2.2 Active Stress Orientation and Anisotropy

In addition to the amount of myosin activity on junctions, tissue stresses depend on its local distribution across junctions. Cortical actomyosin can rigidify an epithelium when isotropically distributed, raising the energy barrier for neighbor exchanges and reducing cell rearrangements [14, 15]. Conversely, anisotropic distributions can drive fluid-like motion, channeling cell rearrangements in certain directions to produce tissue-scale anisotropic flows [14, 15]. The orientation and anisotropy of junctional actomyosin strongly affect the orientation and anisotropy of junctional tension [6, 13] and can sculpt cell shapes [16], orient divisions [17], and direct intercalations [6, 18, 19]. In the avian embryo, the most coherent motifs of myosin anisotropy are supracellular actomyosin cables spanning 2-8 cells, appearing perpendicular to the axis of the incipient primitive streak at the onset of mesoderm contraction [6]. Actomyosin cables are suggested to self-organize in a tension-dependent manner, leading to spontaneous large-scale anisotropies observed in avian gastrulation [4,10,20] and *Drosophila* germband extension [21–24]. To capture the self-organization of actomyosin cables, we treat actomyosin cables as elongated fibers and model their anisotropy (local nematic order) and orientation with a traceless, symmetric tensor 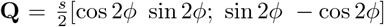; sin 2*ϕ* − cos 2*ϕ*]. Here, 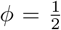 arctan (*Q*_11_*/Q*_12_) represents the average orientation of actomyosin cables and 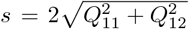 the degree of local cable alignment. Minimally, **Q** evolves due to flow coupling, active alignment, and passive relaxation:

**Supplementary Figure S2:**
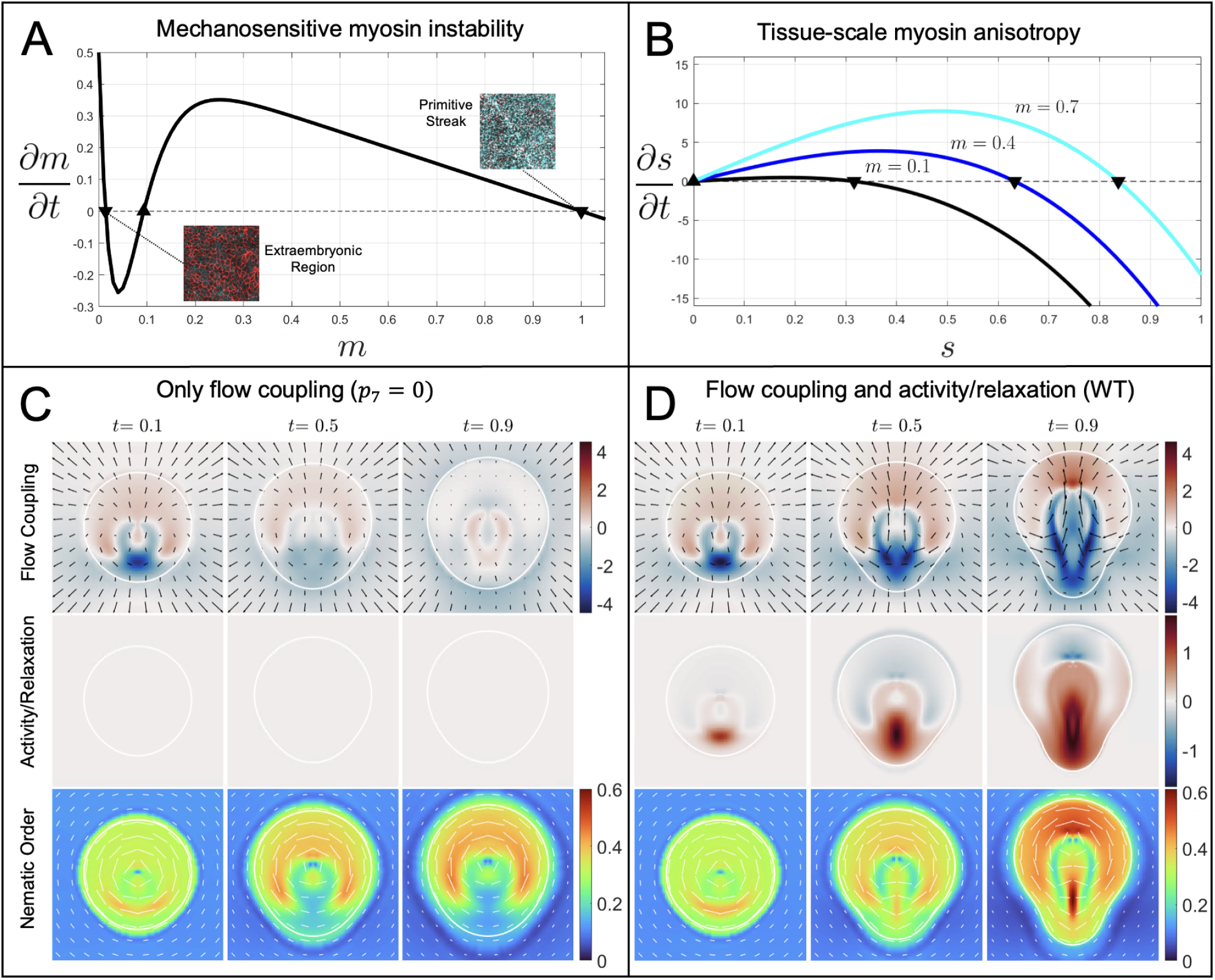
Active Stress Dynamics: A) Spatially uniform equilibria of active myosin (*m*) dynamics (Eq. (S10b)) reveal an unstable fixed point (▲) and a low (high) stable fixed points (▼) towards which EE (EP) regions evolve over time [4]. Insets show actin (red) and doubly phosphorylated myosin light chain (cyan) in the EE and primitive streak regions of an embryo after 15 hours (cf. Fig. S8), depicting tissue states closest to the stable equilibria. B) The flow-free (**v** = **0**) equilibria of active myosin nematic order (*s*) dynamics (Eq. (S5b)) include an instability at *s* = 0 (▲) for *m >* 0 and a higher stable equilibrium (▼) that increases with *m*. C) Without active alignment and relaxation (*p*_7_ = 0), the total derivative of *s* is governed solely by the flow coupling term (first row). The initially elevated nematic order in the posterior is destroyed by the initial convergent extension (third row), and convergent extension does not continue. D) Inclusion of active alignment and relaxation (middle row, *p*_7_ = 25) rescues nematic order from convergent extension, enabling primitive streak formation.

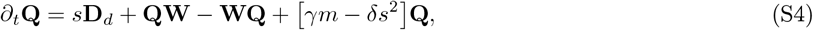

with vorticity tensor 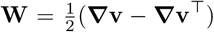 and deviatoric strain rate tensor 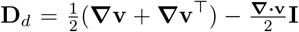. The first term couples cable orientation and anisotropy to the shear rate **D**_*d*_. The second two terms account for rigid body rotation. The final term addresses nematic dynamics independent of tissue flows and only affects *s*, as becomes apparent decomposing Eq. (S4) into the separate dynamics of **Q**’s independent degrees of freedom, *ϕ* and *s* [25]:

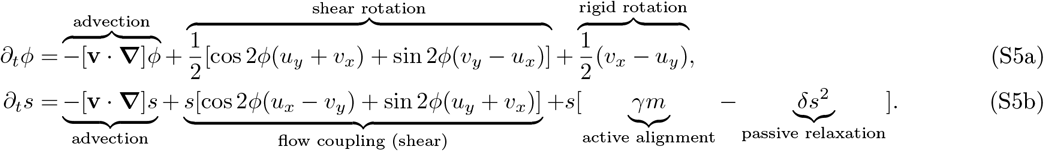

Here, *u*_*x*_, *u*_*y*_, *v*_*x*_, and *v*_*y*_ denote spatial derivatives of *u* and *v* with respect to cartesian coordinates *x* and *y*. We use same notation hereafter for all spatial derivatives. Equation (S5a) is the same as in [4], modelling the passive reorientation of cables by shear and vorticity. Instead, Eq. (S5b) is new and accounts for advection, local alignment or destruction of nematic order by shear, active alignment and passive relaxation. Active alignment (*γ*) in the nematic order dynamics accounts for observations that actomyosin cable formation involves positive feedback, with contracting junctions actively aligning neighboring junctions and anisotropic junctional tension polarizing myosin activity by redistributing myosin to junctions aligned with the direction of highest active stress [6, 10, 13, 16, 26, 27]. Relaxation accounts for the combination of non-directed intercalations and resistance to cell shape deformations, which decrease junctional alignment and favor an isotropic tissue state [14,27]. Without shearing, active alignment will resist relaxation to achieve a nonzero equilibrium nematic order (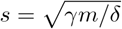, Fig. S2B).

### S2.3 Force Balance

The total tissue stress *σ* includes active (*σ*_*A*_) and viscous (*σ*_*V*_) components:

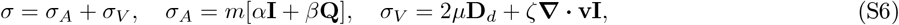

where *µ* and *ζ* denote the shear and bulk viscosities, and *α* and *β* convert *m* to isotropic and anisotropic stresses. The anisotropic active stress term arises from the nematic contractile activity of actomyosin cables. The isotropic active stress term instead represents *i)* the apical constriction of cells by contractile activity of myosin in bulk across a cell’s apical cortex or in bundles along a cell’s perimeter [28] and *ii)* stresses induced by active cell ingressions [4]. Due to the slow flow rates and high viscosity of epithelial dynamics at long times [29], we ignore inertial terms. This leads to an active, compressible Stokes flow (0 = **∇·** *σ*):

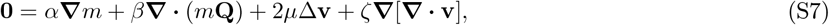

with *x* and *y* components:

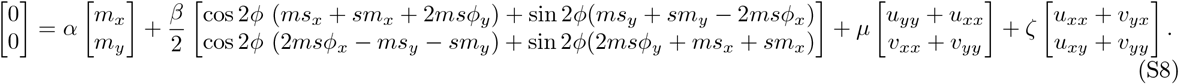

### S2.4 Analysis in 1D and Biophysical Constraints on *α* and *β*

To illustrate the importance of active alignment in Eq. (S5b), we consider a fixed average cable orientation either along (*ϕ*(*x, t*) = 0) or perpendicular to 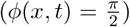 the *x*-axis and uniform in *y*. Neglecting the shear force density 2*µ*Δ**v**, the viscous force density simplifies to *ζ* **∇** [**∇·v**] (*ζu*_*xx*_ in *x*), relating to divergence or convergence along *x*. This reduces the force balance to

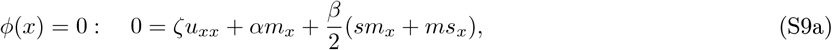

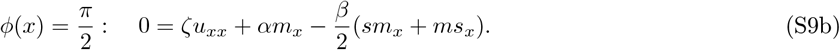

We now ask: given an initial distribution of *s* or *m* with maxima centered at *x* = 0, will a negative velocity divergence be induced at *x* = 0 (as in the primitive streak)? If cables contract along *x* (*ϕ* = 0), a peak in *m* or *s* will unconditionally induce negative divergence at *x* = 0 (Eq. (S9a)), consistent with the cables’ initial tangential orientation resulting in PS formation perpendicular to the cables. However, if cables contract perpendicular to the *x*-axis 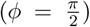, the isotropic (*α*) and anisotropic (*β*) effects of myosin activity compete and their contribution to negative velocity divergence becomes conditional. To explore these conditions, we consider two scenarios for which Eq. (S9b) simplifies:

1. Uniform *m* and locally elevated 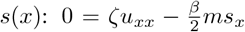. Here, gradients in *s* and *u*_*x*_ are always in the same direction, resulting in no PS formation because divergence becomes more positive as *s* increases.

2. Uniform *s* and locally elevated 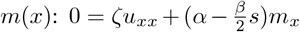. Divergence becomes more negative as *m* increases only if 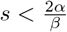 (isotropic active stresses dominate). Since *s* ≤ 1, this condition is always met if 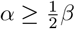.

With our analysis, we choose *β* = *α*, satisfying both of the above conditions and reducing parameters.

The 1D model also offers insights into nematic order dynamics in the avian embryo. For cables aligned perpendicular to the primitive streak (*ϕ*(*x*) = 0), the dynamics of *s* (Eq. (S5b)) becomes 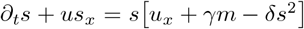. If *s*_*x*_ = 0 (uniform) or *u* = 0 (e.g. in the streak by symmetry), ∂_*t*_*s* = 0 requires 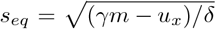. This highlights two competing effects on nematic order: *i)* active cable alignment and *ii)* nematic order destruction by converging flows along the cable orientation, which can be induced by aligned cables. Active alignment can save nematic order if the active alignment rate exceeds any negative divergence induced. These insights explain why, in the 2-dimension wild-type model, convergent extension cannot be sustained with flow coupling alone (Fig. S2C) but can be saved by active alignment (Fig. S2D). To further simplify the model, we set *δ* = *γ*, assuming that alignment and relaxation occur on similar timescales.

### S2.5 Dimensionless 2D Model

We nondimensionalize Eqs. (S3,S4,S7) using a characteristic lengthscale *x*_*c*_ = 4 *mm*, representing the radius of the modeling domain that includes the EP and a fraction of the EE region, and a characteristic timescale *t*_*c*_ = 15 *h*, representing the duration of gastrulation (HH1-HH4). This provides a dimensionless velocity 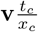 and dimensionless operators *t*_*c*_∂_*t*_, *x*_*c*_ **∇**, and 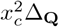. Rearranging terms, we construct a minimal set of dimensionless parameters: *p*_1_ − *p*_7_, defined in Table 1. The full 2D model with dimensionless **v**, *m*, and **Q** becomes

**Table 1:**
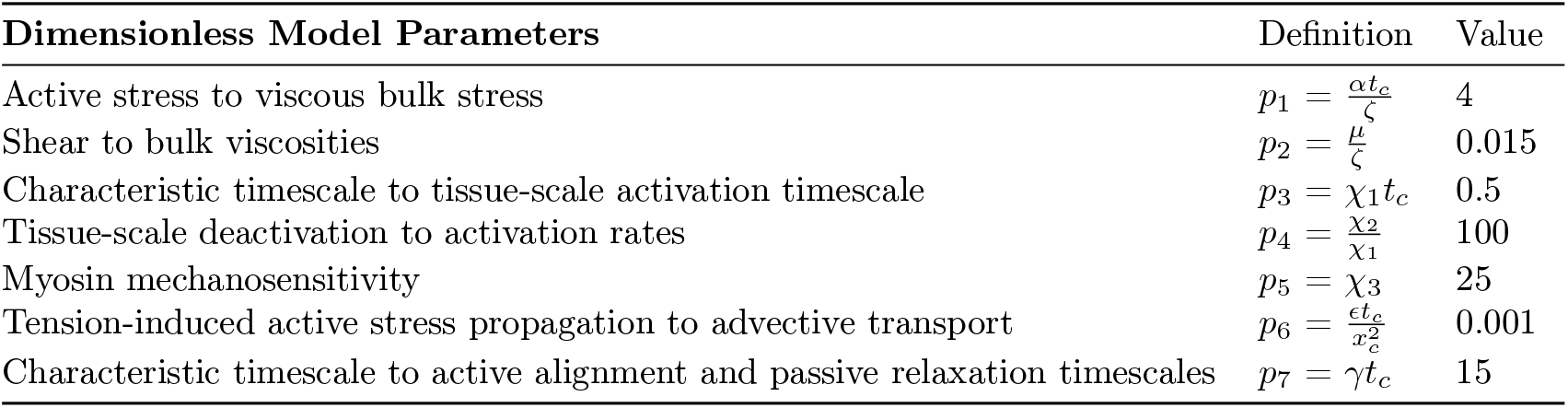
Dimensionless model parameters, their definitions, and default values.

| Dimensionless Model Parameters | Definition | Value |
| --- | --- | --- |
| Active stress to viscous bulk stress | $p_1 = \frac{\alpha t_c}{\zeta}$ | 4 |
| Shear to bulk viscosities | $p_2 = \frac{\mu}{\zeta}$ | 0.015 |
| Characteristic timescale to tissue-scale activation timescale | $p_3 = \chi_1 t_c$ | 0.5 |
| Tissue-scale deactivation to activation rates | $p_4 = \frac{\chi_2}{\chi_1}$ | 100 |
| Myosin mechanosensitivity | $p_5 = \chi_3$ | 25 |
| Tension-induced active stress propagation to advective transport | $p_6 = \frac{\epsilon t_c}{x_c^2}$ | 0.001 |
| Characteristic timescale to active alignment and passive relaxation timescales | $p_7 = \gamma t_c$ | 15 |

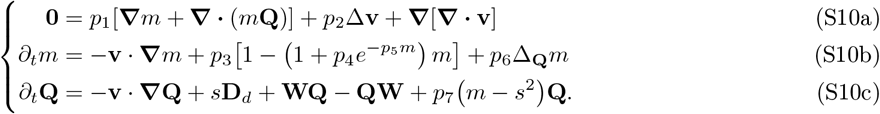

### S2.6 Boundary Conditions

Experiments show a nearly linear initial expansion of the outer extraembryonic radius *R*_*EE*_(*t*) [30, 31], consistent with a constant edge cell crawling speed *v*_*e*_. To model this constant velocity, we use a Dirichlet boundary condition at the outer radius (*R*) of our circular fixed modeling domain: **v**_*R*_(*θ, t*) = [*v*_*R*_(*t*) cos *θ, v*_*R*_(*t*) sin *θ*] (Fig. S3A). Without active stress gradients, constant crawling results in a linearly increasing radial velocity profile *v*_*r*_(*t*) = *v*_*e*_(*r/R*_*EE*_(*t*)) so that *v*_*R*_(*t*) = *v*_*e*_*R/*(*R*_*EE*_(*t*_0_) + *v*_*e*_(*t* − *t*_0_)) (Fig. S3B). While this velocity boundary condition accurately represents epiboly on our fixed modeling domain boundary, our results are robust to changes in *v*_*R*_(*t*) and can also predict observations imposing a constant *v*_*R*_ (Sec. S3). As in [4], the most natural and least intrusive boundary condition for *m* and **Q** is no flux, as experiments suggest no additional sources or sinks of these variables in the EE.

**Supplementary Figure S3:**
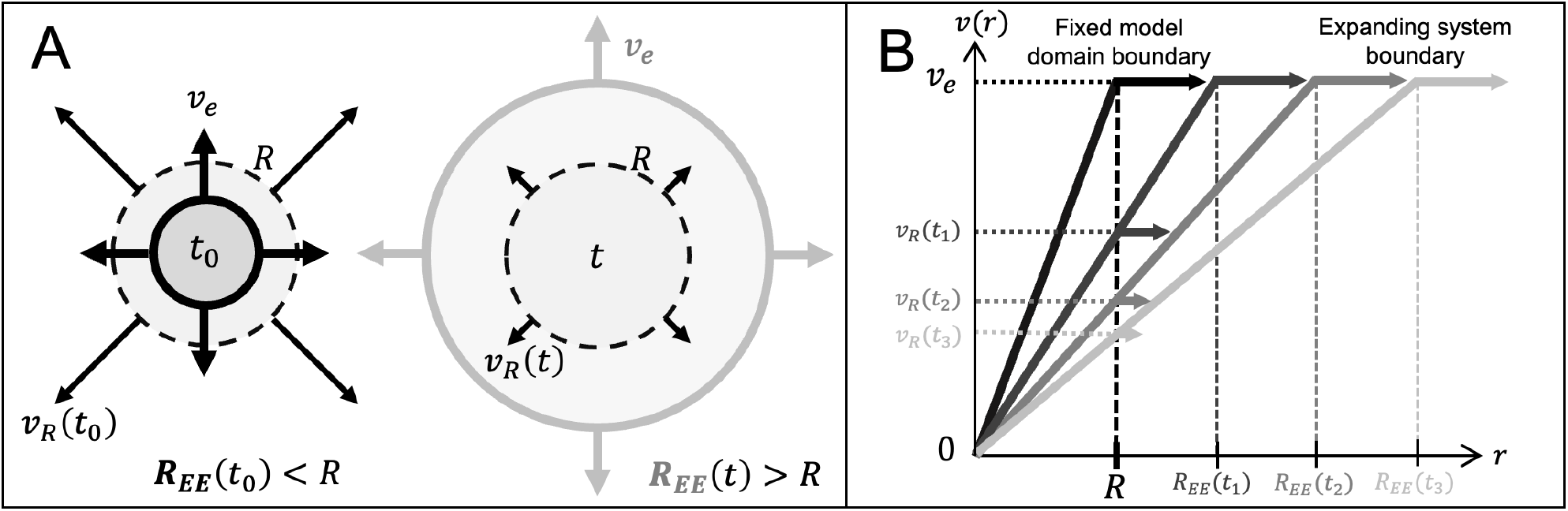
Epiboly-Based Velocity Boundary Condition. A) Outer extraembryonic radius *R*_*EE*_ and model domain radius *R* = *x*_*c*_ at initial and final times. Initial *R*_*EE*_ *< R* ensures that enough extraembryonic material remains in the domain by final time *t* to visualize both repellers. B) Constant outward edge-cell velocity magnitude *v*_*e*_ results in a linear velocity profile *v*(*r*) and therefore a decreasing domain boundary velocity magnitude *v*_*R*_(*t*).

### S2.7 Initial Conditions

Consistent with experiments [4, 6, 13], we model higher initial myosin in the posterior EP with a Gaussian function above a background level *m*_*>*_ = *m*^*^ + 0.001 (Fig. S2A and Fig. 2A), positioned (*r*_*c*_ = 0.25) between the EP center and initial EP boundary (*R*_*EP*_ (*t*_0_) = 0.4). The Gaussian amplitude *A*_*m*_ = 0.25 affects the pace of gastrulation. Its angular and radial extents *σ*_*r,θ*_ =0.03, 0.2 reflect the sickle shape of the presumptive mesoderm at HH1. In the EE, where myosin activity starts and remains low, we initialize myosin *m*_*<*_ below the instability. We set the initial orientation of actomyosin cables *ϕ*(**x**, *t*_0_) along the tangential direction, matching experimental observations [4]. We set the initial nematic order *s*(**x**, *t*_0_) to its equilibrium value in the absence of flow, determined by active alignment, passive relaxation, and the initial distribution of myosin activity (Fig. S2B). This choice is justified by the observed initial anisotropy in cell shapes in the posterior EP, even before the onset of motion [6]. Equation (S10a) determines the velocity at each instant, given *m* and **Q** with boundary velocity **v**_*R*_. We nondimensionalize *r*_*c*_, *σ*_*r*_, *R*_*EP*_ (*t*_0_), and *v*_*e*_ using *x*_*c*_ and *t*_*c*_. The dimensionless effective edge cell velocity *v*_*e*_*/R*_*EE*_(*t*_0_) = 0.5 is consistent with the typical epiboly velocity observed in our experiments (50 − 150 *µm/h*). We use these initial conditions, summarized in Table 2, together with Eq. (S10), to simulate gastrulation over 15 hours (stages HH1-HH4).

**Table 2:** Model initial conditions. Variables are defined in polar coordinates 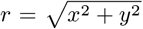 and *θ* = arctan *y/x* computed from the Cartesian coordinates **x** = [*x, y*] on a circular modeling domain centered at **x** = **0**.

| Model Variables | Initial Condition |
| --- | --- |
| Myosin Intensity | $m_0 = \begin{cases} m_> + A_m e^{-\frac{(r-r_c)^2}{2\sigma_r^2} - \frac{(\theta)^2}{2\sigma_\theta^2}} & \text{for } \mathbf{x}_0 \in \mathbf{x}_{EP}, \text{ i.e. } 0 < r \leq R_{EP}(t_0) \\ m_< & \text{for } \mathbf{x}_0 \in \mathbf{x}_{EE}, \text{ i.e. } R_{EP}(t_0) < r \leq R \end{cases}$ |
| Myosin Orientation | $\phi_0 = \theta + \frac{\pi}{2}$ |
| Myosin Anisotropy | $s_0 = \sqrt{m_0}$ |

### S2.8 Numerical Scheme

We solve Eq. (S10) in MATLAB using a finite-difference numerical scheme on a circular domain (Algorithm 1). To compute advection terms, we use first-order upwinding to ensure stability and avoid introducing spurious oscillations. Near the EP-EE boundary, where there are steep gradients in *m* and **Q**, we instead use second-order upwinding to minimize numerical diffusion. We discretize all other spatial differential operators using second-order centered finite differences, except at the boundaries, where we use first-order forward or backward finite differences [32].

#### Algorithm 1 Numerical Solver of Eq. (S10)

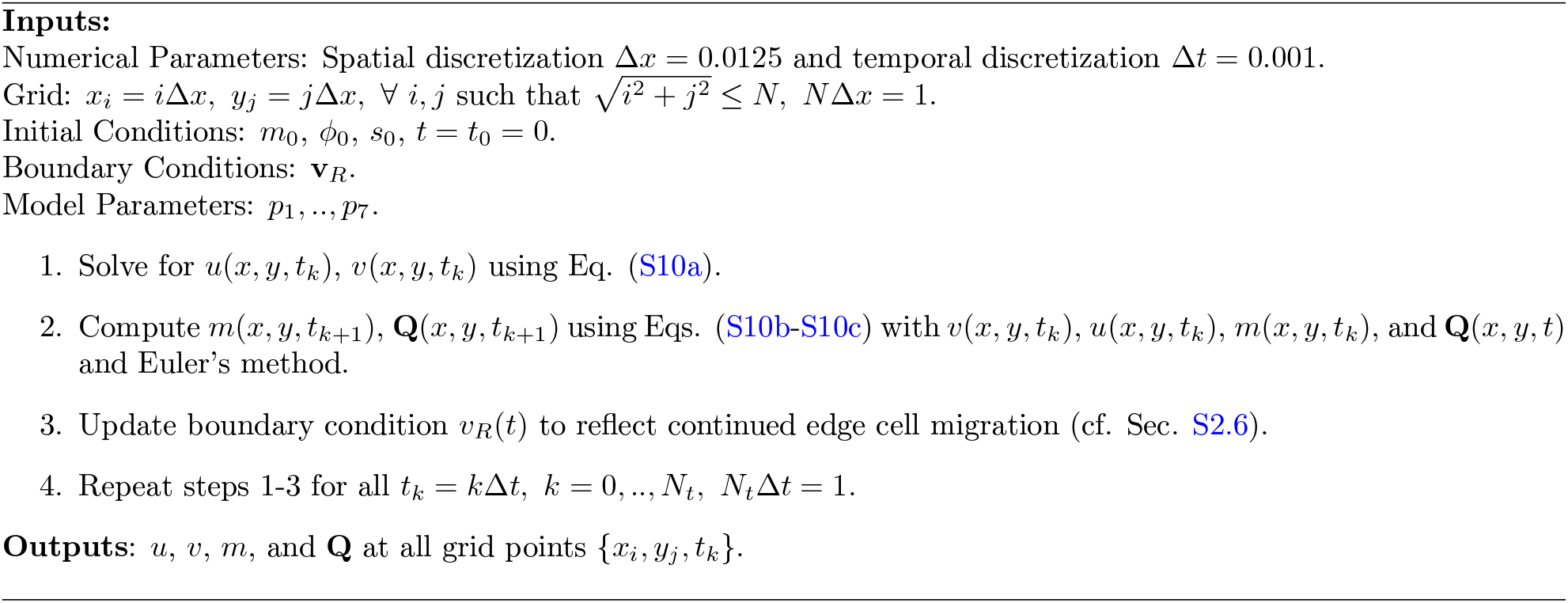

## S3 Model Perturbations

In the main text, we present only model perturbations that are experimentally realizable. To confine the simulated embryo, we blocked epiboly (*v*_*e*_ = 0), resulting in a fixed embryonic radius (*v*_*R*_(*t*) = 0). This change dramatically weakens R1 compared to the wild type. Complete elimination requires eliminating isotropic myosin activity in the EP (Table 3A). Solely eliminating isotropic myosin activity also eliminates R1 (Table 3B), but this is not experimentally realizable without also affecting anisotropic myosin activity [6] (eliminating both repellers). Likewise, the distinct myosin dynamics of the EP and EE are necessary for Repeller 1 (Table 3C). R2 bisects the presumptive mesendoderm region, which, in the model, is characterized by higher initial myosin activity. Accordingly, in the main-text model perturbation, we eliminated the extra posterior myosin (*A*_*m*_ = 0), making the model radially symmetric. While this alone is sufficient to eliminate R2, it can also be achieved by blocking active alignment (*p*_7_ = 0, Table 3D) or anisotropic myosin activity (enforcing *s* = 0, Table 3E). However, these are not feasible in experiments.

**Table 3:**
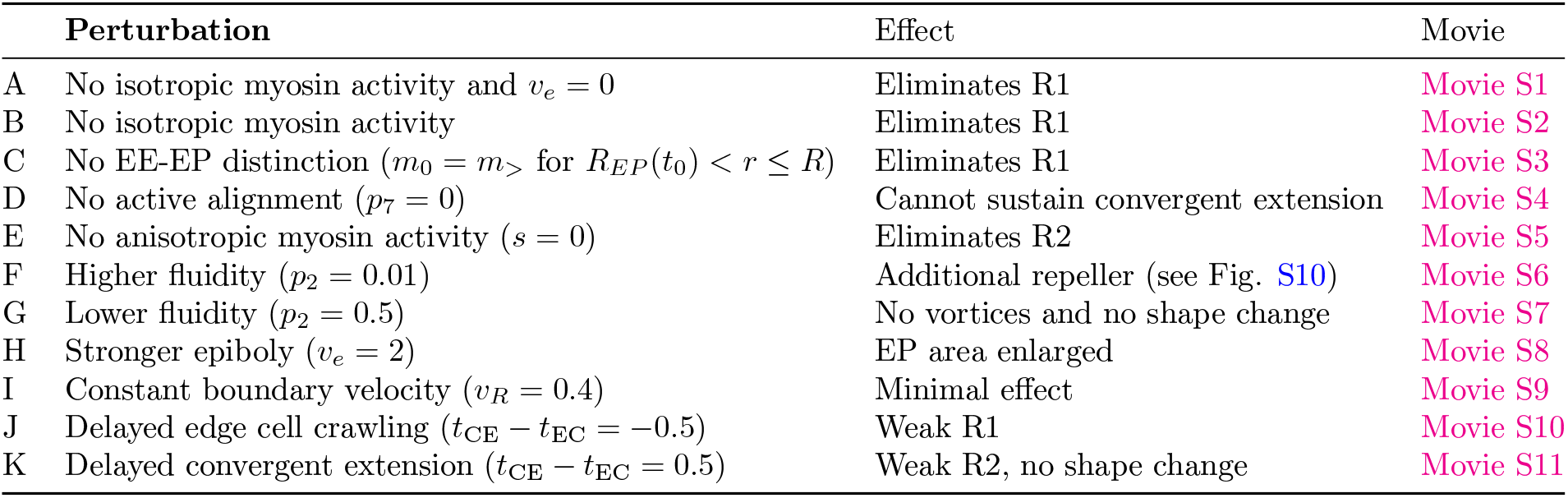
Model perturbations. Associated movies depict the model velocities, velocity divergence, isotropic and anisotropic stresses, active forces, repellers, attractors, and deformed Lagrangian grids over time.

These model perturbations, whose experimental analogs are impractical, help clarify the necessary and sufficient mechanisms for each repeller and their contributions to the embryo’s flows and dynamic geometry. Table 3 includes a series of additional perturbations, varying model parameters, boundary conditions, and initial conditions to elucidate the importance of certain model components and the flexibility of their parameters. Increasing *p*_1_ raises active stress magnitudes and strengthens both repellers (higher 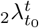). *p*_2_, relating shear and bulk viscosities, encodes tissue fluidity. A more fluidized tissue predicts larger vortices and stronger repellers (Table 3F). Conversely, a less fluid embryo exhibits no large, vortical flows, as previously noted [4], and, crucially, no shape change from circular to pear-shaped (Table 3G). Sensitivity analyses for *p*_3_ − *p*_6_ are studied in [4], where the *m* dynamics is governed by the same biophysical assumptions. Faster edge-cell crawling strengthens R1 and enlarges the EP (Table 3H). Using a constant *v*_*R*_ instead of our dynamic *v*_*R*_(*t*) simplifies the model without substantially affecting our results (Table 3I), but is less realistic.

Finally, we explore the effects of delaying either the onset of epiboly or posterior myosin elevation (Fig. S4). We find that time delays in edge cell crawling weaken R1 while preserving R2 (Table 3J), and time delays in convergent extension weaken R2 while preserving R1 (Table 3K), consistent with their mechanistic modularity. Notably, substantial EP shape change in the model occurs only when convergent extension starts before epiboly, as typically occurs in experiments.

**Supplementary Figure S4:**
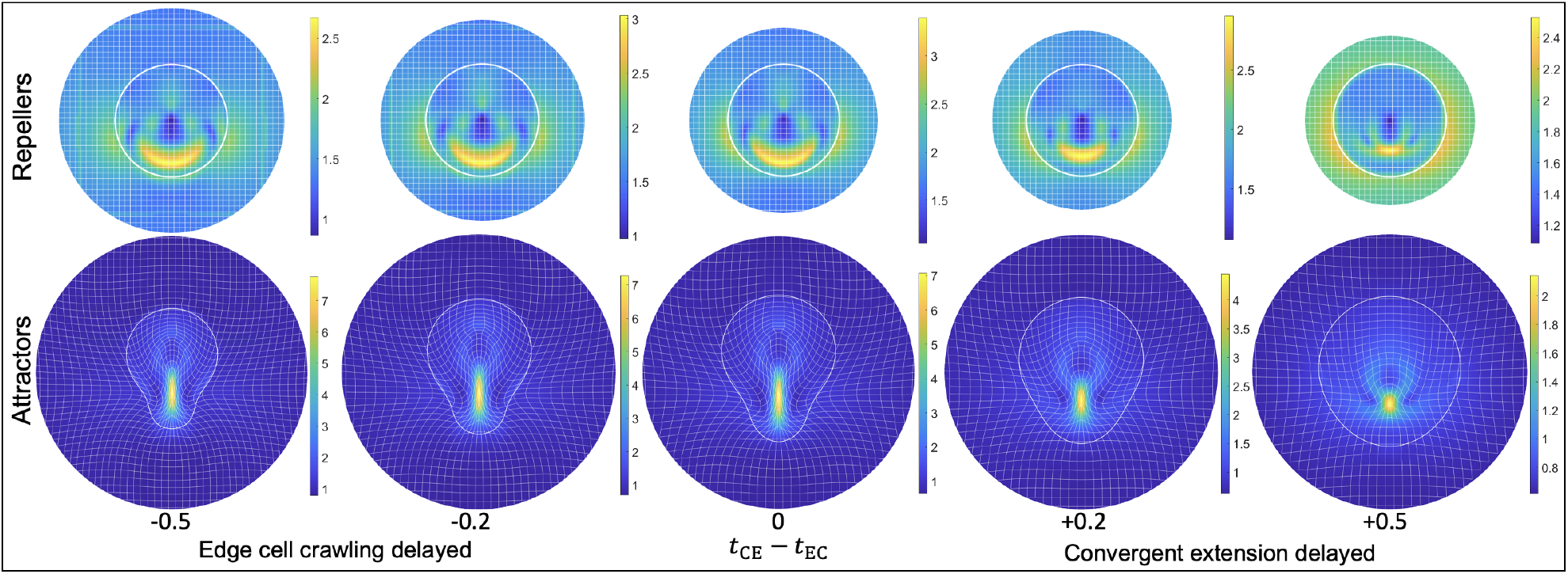
Variable Timing of Epiboly and Posterior Myosin Elevation. Model repellers (attractors) with the initial (final, deformed) Lagrangian grid and EP boundary. *t*_EC_ and *t*_CE_ indicate the time at which edge cell crawling (epiboly) and convergent extension (posterior myosin elevation) start in the model. *t*_0_ is taken to be the smaller of the two. Columns depict output for different values of *t*_CE_ − *t*_EC_. If *t*_EC_ *> t*_CE_, posterior myosin is initially elevated but *v*_*e*_ = 0 until *t* = *t*_EC_ (left two columns). If *t*_EC_ *< t*_CE_, EP myosin is initially uniform, but we add posterior myosin (Gaussian function in Table 2) at *t* = *t*_CE_ (right two columns). In the middle column, *t*_CE_ = *t*_EC_ = *t*_0_. Colormaps show the unnormalized _2_*λ* fields.

### S3.1 Active Forces

Movies 1-4 and S1-S11 include the magnitude and direction of active forces *p*_1_[**∇** *m* + **∇·** (*m***Q**)] (cf. Eq. (1a)), indicating, in addition to the boundary velocity, the active drivers of the gastrulation flows. Active forces are strong around the presumptive mesendoderm and later in the vicinity of the developing streak consistent with our earlier work [4]. These active forces, arising from the active intercalation and ingression of mesendoderm cells, generate EP vertical flows, convergent extension and shape change. In addition, the gradient in myosin activity between the EE and EP results in inward active forces constricting the EP at its boundary. These active forces largely depend on isotropic myosin activity and generate Repeller 1 (Table 3B) but are insufficient to drive gastrulation flows and embryo shape change alone (Table 3E). In fact, removing the differential EE-EP myosin dynamics eliminates this second pattern of active forces (Table 3C).

## S4 Note on Epiboly and Evolution

Changes in the amount of yolk in the egg have been recognized as one of the main drivers of evolutionary change during gastrulation [33–35]. Amniotes (reptiles, birds, and mammals) evolved from ancestors that could lay eggs on land, achieved through the acquisition of a mineralized shell that provided mechanical protection and three extraembryonic membranes [36]. Most importantly, the yolk experienced a significant increase in size, probably to increase the egg’s energy depot, allowing the embryo to hatch as a miniature version of the adult instead of as a larva [37]. This increase in yolk size generated mechanical and topological constraints that required adaptations in the gastrulation mode.

The early cleavages became exclusively meroblastic due to the impossibility of splitting the enormous yolk cell [36,38], leading to the formation of discoidal embryos that sit on top of the yolk. This topological configuration imposes a problem on the developing embryo, as it now has to engulf the yolk underneath to effectively access its nutrients. Consequently, large yolks are associated with the evolution of extraembryonic tissues that engulf the yolk and later help to digest it, as seen in teleost fish and amniotes. The main role of epiboly, the process by which extraembryonic tissues expand and thin to enclose the yolk, is to form the yolk sac and provide nutrients to the embryo later in development [39].

In amniotes, including chicken, the extraembryonic tissues have acquired an additional role in patterning the embryo during gastrulation. For example, in chick, FGF8 and WNT9C secreted by the the hypoblast and the EE help to position the mesoderm [40]. However, the importance of the mechanical inputs imposed by the EE on the embryo proper (EP) and their significance for development remains an open question. Epiboly induces global tension [30, 31, 39, 41] that propagates to the EP, as severing the EP-EE boundary causes both regions to contract [39]. Previous studies have suggested that epiboly is necessary for correct early development in avian embryos, as ablating large EE areas at primitive streak stages leads to poor embryo development [39]. However, it is unclear if this is due to the removal of mechanical inputs or the disruption of signaling roles played by the EE.

Our findings reveal that epiboly is not required for the early stages of avian embryogenesis. When we confined the embryo to eliminate epiboly while maintaining an intact EE, embryos still gastrulated and developed complete axial structures (brain, somites, neural tube, tailbud), despite having shorter, but proportioned, body axes (Fig. S9). Recent reports have shown that increasing tension in the chick embryo results in a shorter body axis [42]. Similarly, disrupting epiboly progression in zebrafish does not prevent gastrulation but produces a shorter body axis [43, 44]. These findings indicate that while epiboly movements are not required for gastrulation to occur, they can impact embryo body length at later stages.

This suggests that the evolution of the amniote egg likely necessitated the development of mechanisms to maintain the shape and integrity of the EP during epiboly. Embryo contractility in amniotes may have evolved as a mechanism to resist the influence of epiboly, allowing the embryo to maintain its intrinsic patterning mechanisms and developmental timeline, ensuring proper body plan formation. This idea is supported by experiments showing that partial EP ablations normally require detachment of the EE from the vitelline membrane to avoid the embryo ripping apart [45], suggesting that epiboly forces can pull apart a mechanically compromised EP.

The evolution of the amniote egg, with its increased yolk size and extraembryonic membranes, has led to significant adaptations in the gastrulation process. While epiboly is essential to construct the structure that will provide nutrients to the developing embryo, our findings suggest that it may not be strictly required for the early stages of avian embryogenesis. In fact, epiboly might even be a developmental burden that the early embryo must overcome. The contractility of the amniote embryo may have evolved as a mechanism to maintain its shape and integrity during epiboly, allowing it to follow its intrinsic developmental program without being excessively influenced by the mechanical forces imposed by the extraembryonic tissues. These insights underscore the complex interplay between mechanical forces, signaling pathways, and evolutionary adaptations in shaping the gastrulation process and embryonic development.

## S5 Experimental Methods

### S5.1 *Ex Ovo* Culture

Fertilised white Leghorn chicken eggs were incubated at 37°C for 1-6 hours to around Hamburger Hamilton stage 1 (HH1). Embryos were isolated on a windowed piece of filter paper and cultured *ex ovo* on a semisolid agar-albumen medium, ventral side up [46]. To acquire high-quality velocity fields capturing cell movements on the embryo surface, it is essential to remove as much yolk as possible from the embryos. Embryos were cleaned with a mini Pasteur pipette using a warm 0.1% Tween in physiological saline solution and rinsed with physiological saline solution. For high-throughput live imaging, a large transparent plate was covered with semisolid agar-albumen medium surrounded by wet paper and in the presence of antibiotics, where the embryos were arranged in a grid (Fig. 5A) with the ventral side up.

**Supplementary Figure S5:**
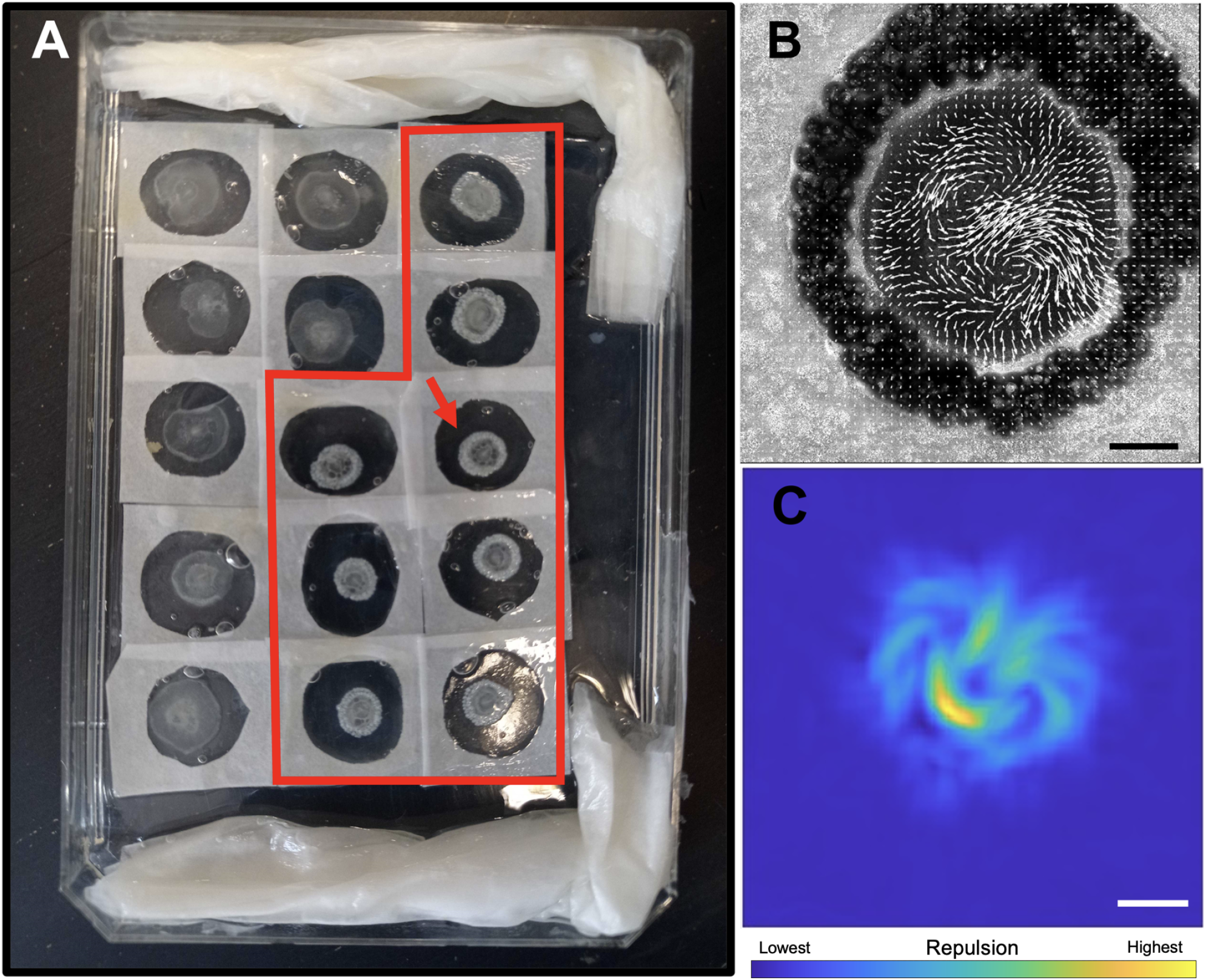
High Throughput Quantification of Chick Gastrulation Flows. A) Plate of 15 chick embryos culture *ex-ovo* for automatic wide-field time-lapse imaging using a motorized stage. The red polygon delineates confined chick embryos. The arrow indicates the embryo shown in B-E. B) Quantification of tissue flows using PIV. Arrows are enlarged for visualization. C) Large values of 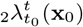 computed from PIV velocities mark repellers (cf. Sec. S1). Scale bars are 1 mm. Movie S12 shows the simultaneous development of 8 wild type (top) and 8 confined (bottom) embryos. Each box containing an embryo is 8.3 mm wide.

### S5.2 Mechanical Confinement and Chemical Perturbations

To mechanically confine embryos we cultured chick embryos on a piece of filter paper with a wide opening (∼15 mm). Later, we cauterized the embryonic side of the vitelline membrane surrounding the embryo using a soldering iron at 250°C, creating a fixed boundary of denatured material that prevents epibolic expansion. For chemical perturbations, a pan-FGF receptor inhibitor (LY2874455, 1*µM*, SelleckChem) was added to the culture medium to block mesoderm differentiation as in [47]. Control embryos were cultured under identical conditions without confinement or in the presence of 0.1% DMSO.

### S5.3 Velocimetry

Embryos were imaged with a Nikon Eclipse Ti inverted bright-field microscope equipped with a motorized stage, a 20X objective, and an Orca Flash 4.0 camera. Tile scan images (6×6 or 7×7) were acquired every 30 minutes for up to 24 hours. When using chemical inhibitors, the embryos were imaged in a 6-well plate with individual agar-albumen substrates for each embryo to avoid chemical diffusion to control samples. Velocity fields were computed from timelapse movies of developing embryos using PIVLab v2.56 for MATLAB with three passes of 576×576, 144×144 and 72×72 pixel interrogation windows with 50% overlap with default preand post-processing parameters. Deformation grids, decomposed strain rates, and the DM were then computed from the velocity field as in [1, 6, 47, 48].

### S5.4 Determination of Embryo Proper, and Extraembryonic Areas Over Time

To determine the embryo size over time, a polygon is manually drawn around the embryo’s edge and tracked using the velocity fields. To obtain the the area of the EP over time the repeller field of the DM 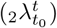 is overlapped on the embryo as a guide to draw a polygon at the EP boundary.

### S5.5 Immunohistochemistry

Embryos were fixed in 4% paraformaldehyde overnight at 4°C, permeabilized with PBS 0.1% Tween (PBT), and blocked with 10% goat serum and 2% bovine serum albumin in PBT. A primary rabbit antibody against double phosphorylated Thr-18 / Ser-19 and myosin light chain 2 (3674, Cell Signaling Technology) was applied overnight at 4°C. After washing, embryos were incubated with an Alexa Fluor™ 555 goat anti-rabbit secondary antibody (Invitrogen) in the presence of Alexa Fluor™ Plus 405 Phalloidin (Invitrogen) and SYTOX™ Deep Red Nucleic Acid Stain (Invitrogen) overnight at 4°C. Finally, embryos were washed, mounted between two thickness 1 cover slides with VECTASHIELD® Antifade Mounting Medium (Vector Laboratories), and imaged using a Zeiss LSM700 confocal microscope with a 20X or a 40X objective and 0.5 zoom.

### S5.6 Computational Surface Extraction

Active myosin resides in the most apical compartment of the cells. For this reason, maximum projections may not accurately represent the apical myosin, including the myosin cables, across the embryo. To tackle this problem we computationally acquire a 2D surface of the most apical side of the embryo. To extract the apical surface of the embryo from the confocal image volumes, we used a custom MATLAB script based on the square gradient focusing algorithm [48, 49]. The steps are as follows:

1. Tile the confocal volume into columns.
2. Apply the square gradient focusing algorithm to each column to find the surface position.
3. Generate a height map by finding the depth of the fastest change in the image sharpness (peak of the second derivative) for each column.
4. Apply a smoothing filter to the height map.
5. Use the smoothed height map to section the image volume and produce a 2D image of the embryo surface.

## S6 Additional Supplementary Figures

**Supplementary Figure S6:**
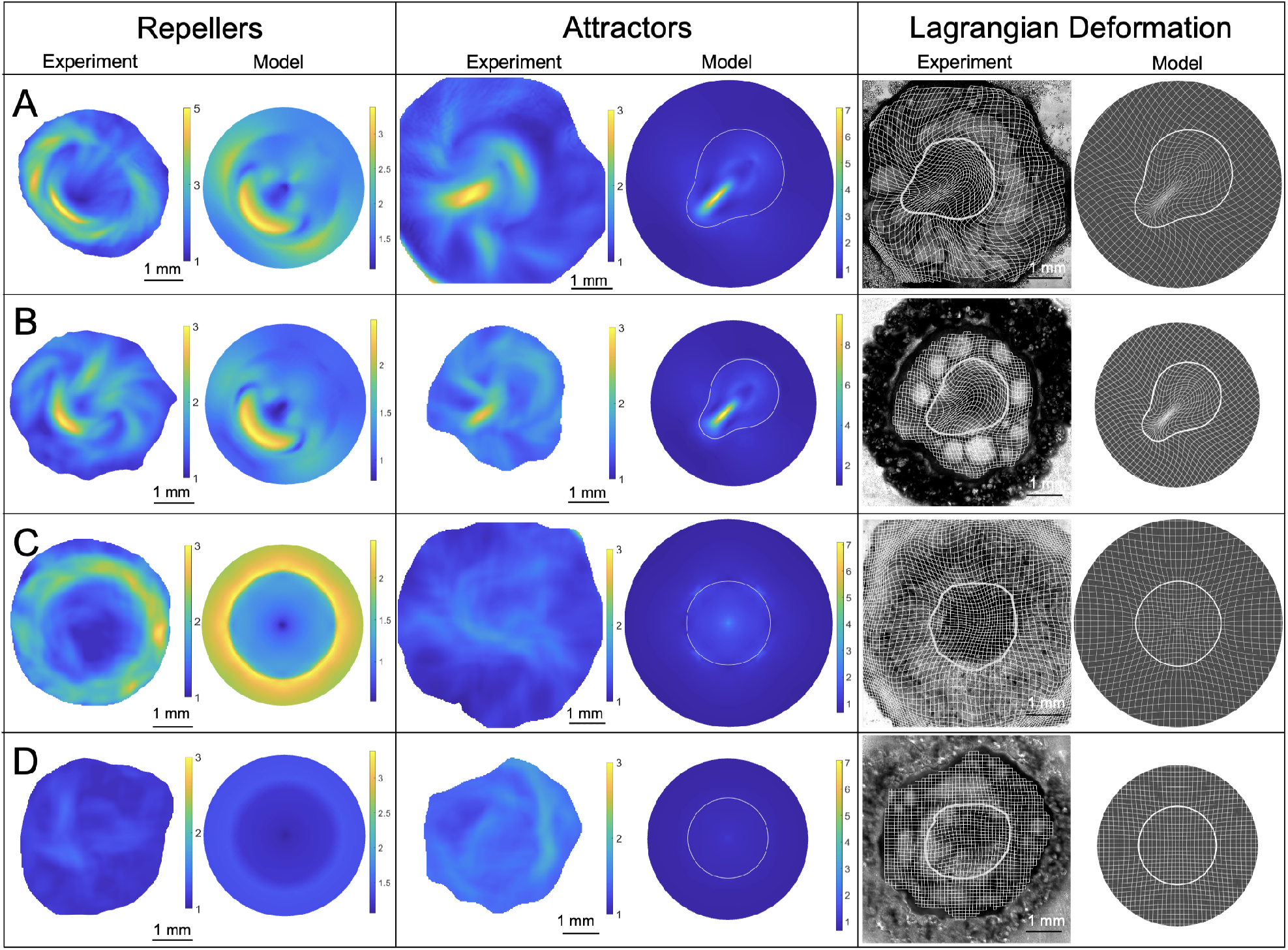
Dynamic Morphoskeleton in Main-Text Perturbations. Repellers, Attractors and Lagrangian deformation for combinatorial elimination of R1 and R2. The attractor marks the PS (cf. Fig. 1). White curves mark the EP boundary at final times *t*. A) Wild type, both repellers present. B) Confined, R1 eliminated. C) No mesoderm, R2 eliminated. D) Combined, both repellers eliminated. Eliminating R2 eliminates the attractor. Colorbars for repellers (attractors) mark un-normalized values of 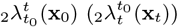. The repeller field in C and repeller and attractor fields in D use the WT colorbar (A) to emphasize the relative lack of deformation.

**Supplementary Figure S7:**
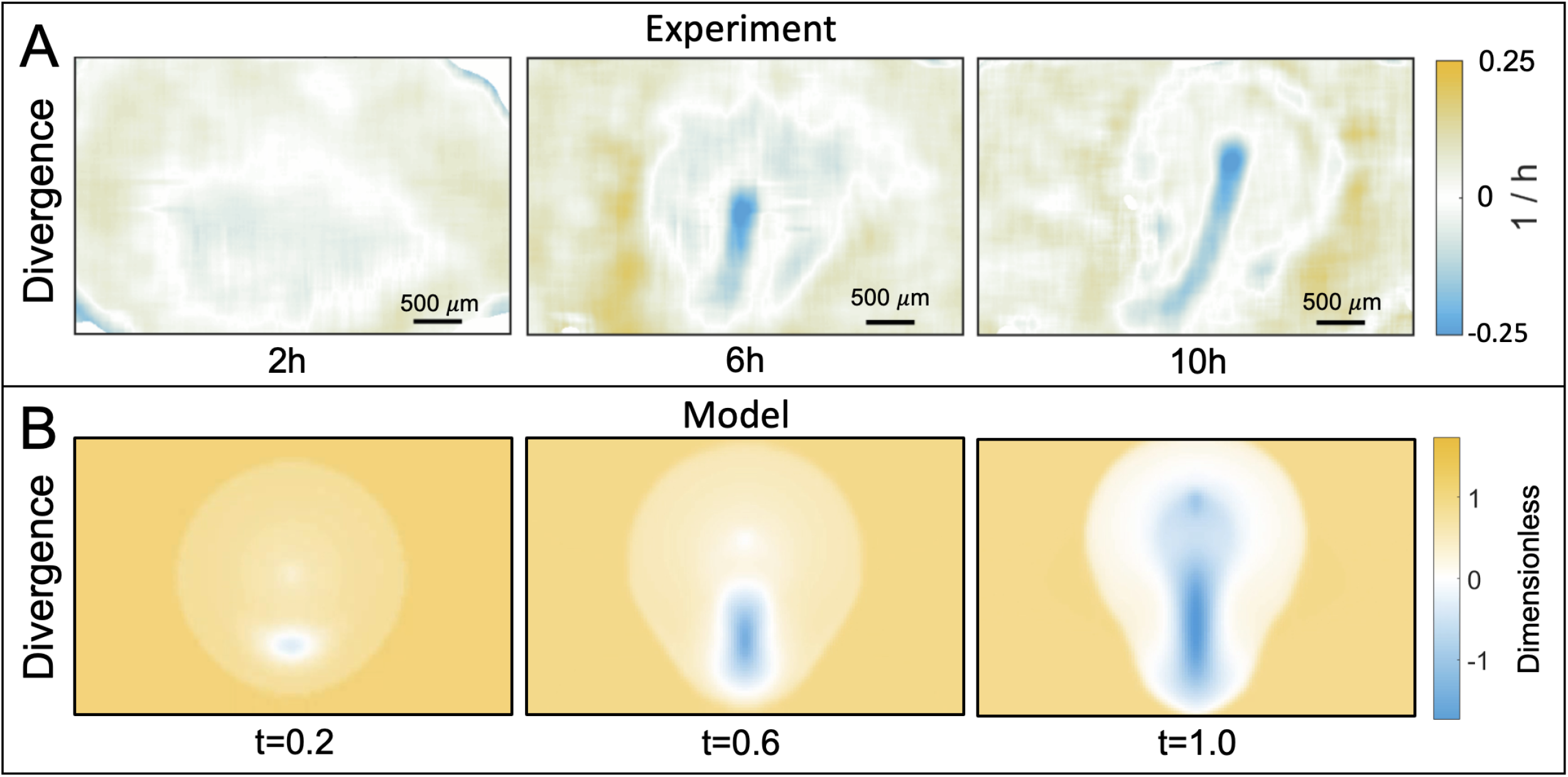
Velocity Divergence. A) Velocity divergence **∇· v** in wild-type experiments (A) and model (B) at three times. In A, 0 *h* corresponds to HH1.

**Supplementary Figure S8:**
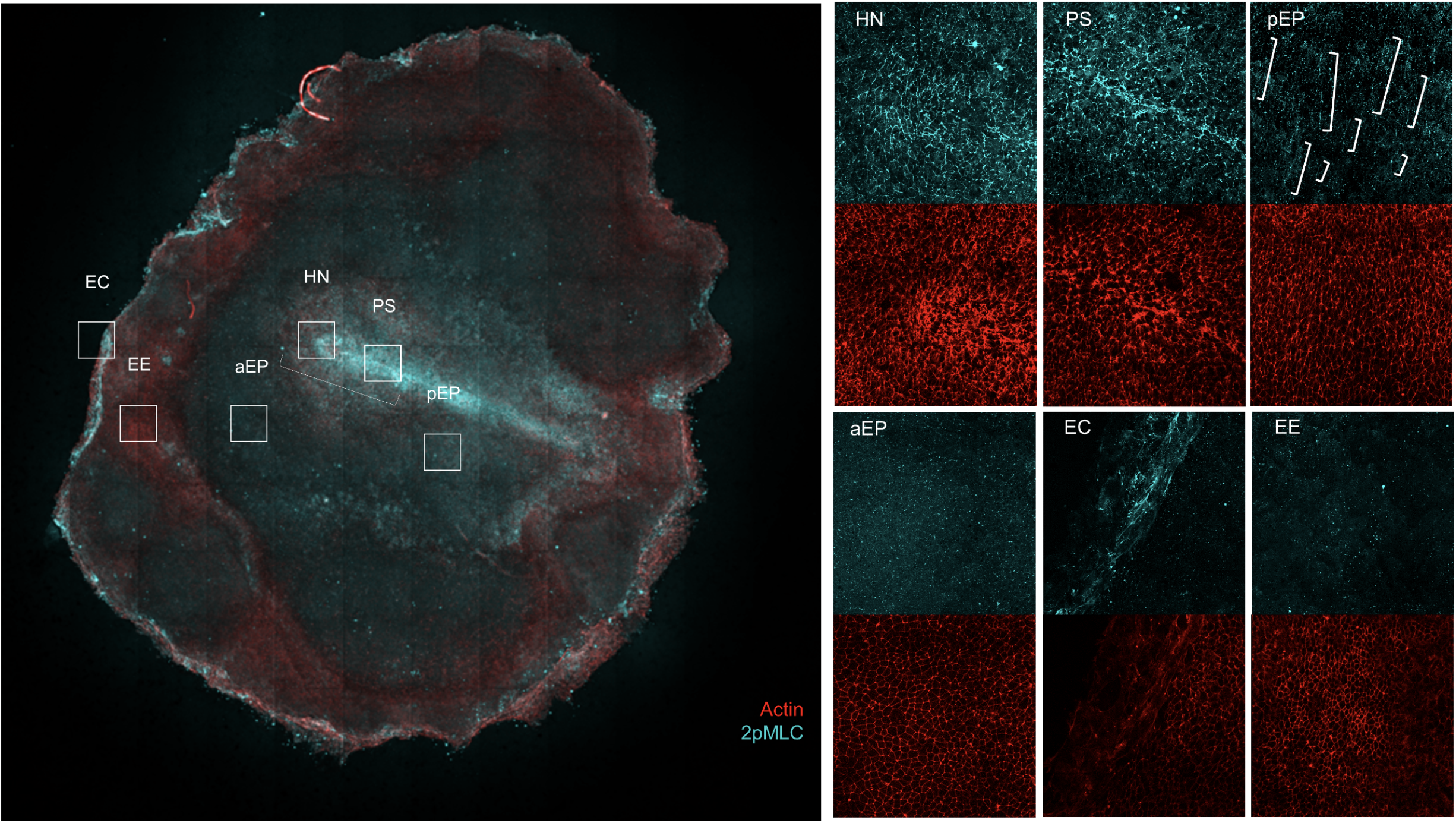
Characterisation of Myosin Patterns in a Confined Embryo. Overview) Full image of a confined embryo at stage HH3+. Each box is 400 *µm* in side. HN) Hensen’s node, PS) Primitive streak, pEP) Posterior embryo proper, note long myosin cables perpendicular to the PS aEP) Anterior Embryo proper, EC) Edge cells, EE) Extraembryonic territory.

**Supplementary Figure S9:**
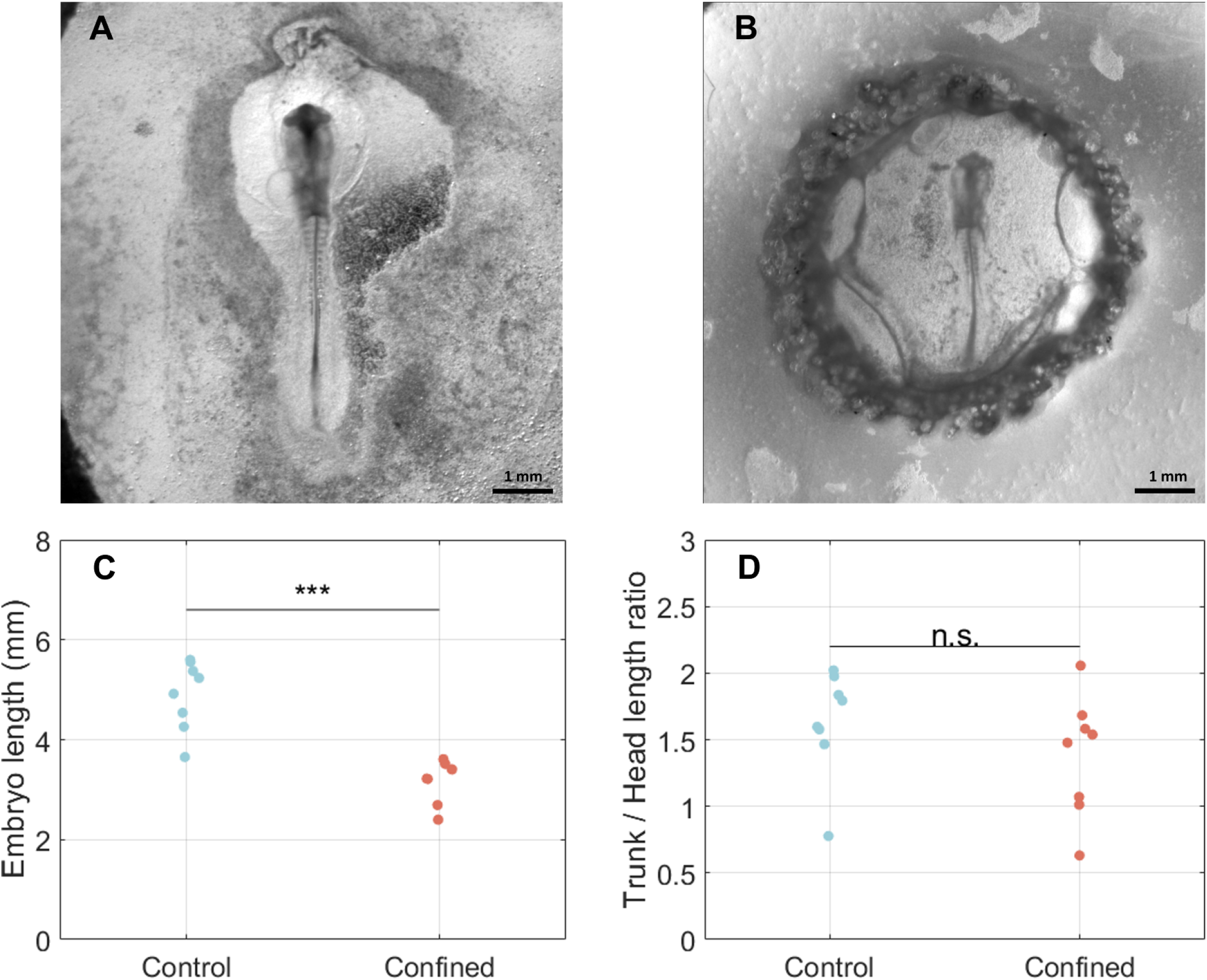
Embryos Are Shorter but Proportional After 48 Hours of Confined Development. A) Embryo after 48 hours of *ex-ovo* development. B) Confined embryo after 48 hours of *ex-ovo* development. C) Embryo length for control and confined embryos after 48 hours of development. D) Trunk/head length ratio for control and confined embryos after 48 hours of development. N=8, Statistical analysis: Two-sample t-test, p-value<0.001 (***), p-value>0.05 (not-significant, n.s.).

**Supplementary Figure S10:**
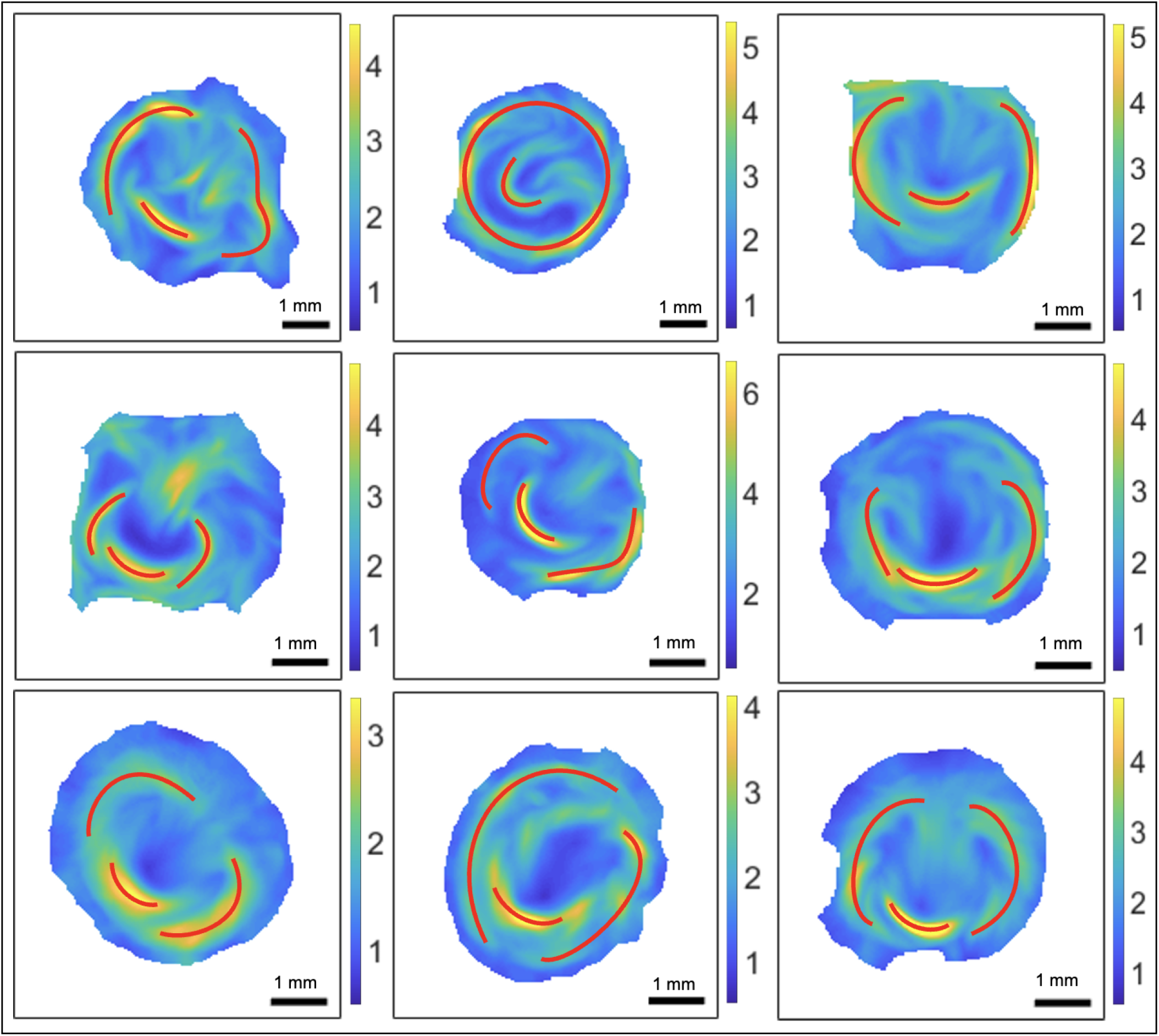
Robustness and intrinsic Variability of Repellers. Experimental unnormalized 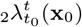 for 9 wild-type embryos recorded for ≈15 *h*. Red curves mark the repellers (R1 and R2). Each repeller’s precise shape and position vary, but R1 consistently wraps around the initial EP boundary while R2 consistently arcs across the posterior. We note the variability in the intensities of R1 and R2 as well as the occasional appearance of a third repelling structure in the anterior, associated with lateral separation (perpendicular to the AP axis) of cells at opposite sides of the PS when it approaches the anterior EP boundary. This feature is also present in the model repeller field (Fig. 2G) and visualizable from the co-located tangential deformation in the Lagrangian grid (Fig. 2F). Note the robustness of Repellers and the DM despite our high throughput experimental approach using a bright-field microscope (Sec. S5) generating less resolved tissue flows compared to single-embryo experiments from a light-sheet microscope (cf. Fig. 3A in [1]).

## Supplementary movies

Movie 1: Time evolution movie associated with Fig. 2.

Movie 2: Time evolution movie associated with Fig. 3.

Movie 3: Time evolution movie associated with Fig. 4.

Movie 4: Time evolution movie associated with Fig. 5.

Movie S1: Time evolution of the fields in Movie 1 for perturbation with no isotropic myosin activity and no epiboly (*v*_*e*_ = 0).

Movie S2: Time evolution of the fields in Movie 1 for perturbation with no isotropic myosin activity.

Movie S3: Time evolution of the fields in Movie 1 for perturbation with no EE-EP distinction (*m*_0_(**x**_*EE*_) = *m*_*>*_).

Movie S4: Time evolution of the fields in Movie 1 for perturbation with no active alignment or passive relaxation (*p*_7_ = 0).

Movie S5: Time evolution of the fields in Movie 1 for perturbation with no anisotropic myosin activity (*s* = 0).

Movie S6: Time evolution of the fields in Movie 1 for perturbation with higher fluidity (*p*_2_ = 0.01).

Movie S7: Time evolution of the fields in Movie 1 for perturbation with lower fluidity (*p*_2_ = 0.5).

Movie S8: Time evolution of the fields in Movie 1 for perturbation with stronger epiboly (*v*_*e*_ = 2).

Movie S9: Time evolution of the fields in Movie 1 for perturbation with a constant domain boundary velocity *v*_*R*_ =constant (0.4).

Movie S10: Time evolution of the fields in Movie 1 for perturbation with delayed edge cell crawling (*t*_*CE*_ − *t*_*EC*_ = − 0.5), corresponding to the leftmost column of Fig. S4.

Movie S11: Time evolution of the fields in Movie 1 for perturbation with delayed convergent extension (*t*_*CE*_ − *t*_*EC*_ = 0.5), corresponding to the rightmost column of Fig. S4.

Movie S12: Time evolution of bright-field microscope images of 8 wild-type (top) and 8 confined (bottom) embryos. Each box containing an embryo is 8.3 mm wide.

